# A High-Throughput Pipeline Identifies Robust Connectomes But Troublesome Variability

**DOI:** 10.1101/188706

**Authors:** Gregory Kiar, Eric W. Bridgeford, William R. Gray Roncal, Consortium for Reliability and Reproducibility (CoRR), Vikram Chandrashekhar, Disa Mhembere, Sephira Ryman, Xi-Nian Zuo, Daniel S. Margulies, R. Cameron Craddock, Carey E. Priebe, Rex Jung, Vince D. Calhoun, Brian Caffo, Randal Burns, Michael P. Milham, Joshua T. Vogelstein

## Abstract

Modern scientific discovery depends on collecting large heterogeneous datasets with many sources of variability, and applying domain-specific pipelines from which one can draw insight or clinical utility. For example, macroscale connectomics studies require complex pipelines to process raw functional or diffusion data and estimate connectomes. Individual studies tend to customize pipelines to their needs, raising concerns about their reproducibility, and adding to a longer list of factors that may differ across studies (including sampling, experimental design, and data acquisition protocols), resulting in failures to replicate. Mitigating these issues requires multi-study datasets and the development of pipelines that can be applied across them. We developed NeuroData’s MRI to Graphs (NDMG) pipeline using several functional and diffusion studies, including the Consortium for Reliability and Reproducibility, to estimate connectomes. Without any manual intervention or parameter tuning, NDMG ran on 25 different studies (≈ 6,000 scans) from 15 sites, with each scan resulting in a biologically plausible connectome (as assessed by multiple quality assurance metrics at each processing stage). For each study, the connectomes from NDMG are more similar within than across individuals, indicating that NDMG is preserving biological variability. Moreover, the connectomes exhibit near perfect consistency for certain connectional properties across every scan, individual, study, site, and modality; these include stronger ipsilateral than contralateral connections and stronger homotopic than heterotopic connections. Yet, the magnitude of the differences varied across individuals and studies—much more so when pooling data across sites, even after controlling for study, site, and basic demographic variables (i.e., age, sex, and ethnicity). This indicates that other experimental variables (possibly those not measured or reported) are contributing to this variability, which if not accounted for can limit the value of aggregate datasets, as well as expectations regarding the accuracy of findings and likelihood of replication. We, therefore, provide a set of principles to guide the development of pipelines capable of pooling data across studies while maintaining biological variability and minimizing measurement error. This open science approach provides us with an opportunity to understand and eventually mitigate spurious results for both past and future studies.

## 1. Introduction

Recent developments in technology enable experimentalists to collect increasingly large, complex, and heterogeneous data. Any single study can include both raw multi-modal data and extensive metadata, including sample design, experimental protocols, data acquisition, and subject specific demographics/phenotypics. Each of these variables adds different sources of variability, which can hamper our ability to interpret and/or generalize results [1; 2]. Moreover, often only a subset of the potential sources of variability are documented or reported [3]. Interpreting these data therefore requires deep data processing pipelines. Such pipelines are particularly important when attempting to enlarge sample size and increase power by aggregating data across multiple studies. These pipelines, however, can introduce additional sources of variability, if different pipelines are used on different datasets, or the same pipeline is applied across datasets but requires substantial tuning or manual intervention, or is run using different operating systems [4]. These sources of variability can collectively swamp the signal of interest, yielding studies with questionable reproducibility, scientific validity, and/or clinical utility.

Studies in neuroimaging exemplify these properties. The data from a single study consist of structural, functional, and/or diffusion magnetic resonance imaging (sMRI, fMRI, and dMRI) scans, from multiple individuals. The metadata associated with a study includes the time, date, and location of the scans, the make and model of the scanner hardware and software, scanner acquisition protocols, as well as the demographic information from each individual, only some of which may be recorded and reported. A number of investigators have developed processing pipelines for one or more of these modalities [5–16], but none of the pipelines are designed to address the variability alluded to above, or work across both fMRI and dMRI.

For example, a number of studies have identified frequently uncontrolled variables that can radically alter the results of downstream inferences, such as menstrual cycle status [3]. Others have demonstrated that some statistics and normalization procedures result in stable parameter estimates across fMRI measurements *within* an individual and study, but those analyses lacked a coherent statistical model, did not compare across studies, and did not consider dMRI data [3]. A few investigations have pooled data across studies, with mixed results. With enough samples, certain properties are apparently preserved [17; 18]. Alternately, the use of sophisticated machine learning techniques can mitigate some of these issues [19]. Nonetheless, many studies continue to fail to be replicated [20].

To rigorously identify and quantify the sources of variability both within and across multi-modal neuroimaging requires (1) data and (2) a pipeline. The requisite data includes numerous datasets with multiple measurements per individual–including data that both conserve and vary a number of different factors. The requisite pipeline must be able to fully process each sample and study, and analyze the results both within and across studies using a coherent statistical model.

The Consortium of Reliability and Reproducibility (CoRR) consists of about 30 different studies from nearly 20 different institutions around the world that provide the necessary data [21]. But no existing pipeline can successfully estimate and meaningfully process every scan in this dataset–including both functional and diffusion data—while also quantifying the magnitude and source of variability amongst them. We, therefore, established several principles and metrics to guide the development of pipelines. We developed an approach, “Neuro Data MRI to Graphs” (NDMG), that meets or exceeds standards along each of the above mentioned principles. We validated our pipeline by running NDMG on 11 dMRI studies comprising 3,227 individuals with 4,347 scans, and 18 fMRI studies comprising 714 individuals with 1,646 scans. For each scan NDMG estimates a “connectome” (a functional or structural map of connectivity) at 24 different spatial resolutions—yielding a total of > 100, 000 estimated connectomes—all of which are publicly available from http://m2g.io. This is the largest open database of connectomes [22], and one of the largest mega-analyses (inference across studies) of multi-modal connectomics data to date [19; 23].

These connectomes provided the data that led us to develop statistical connectomics methods to quantify various connectome properties, such as the relative probability of ipsilateral vs. contralateral connections and homotopic vs. heterotopic connections. While these properties have been previously noted in single studies [24–26], this work demonstrates that aspects of these properties are preserved both across individuals and studies upon optimizing and harmonizing the pipeline. Nonetheless, within session, site, study, and demographic cohorts, substantial variability remained in the *magnitude* of these properties. Moreover, we observed a considerably higher degree of variability across sites, studies, and demographic cohorts. In part, this variability may be due to legitimate biological heterogeneity that could not be accounted for using the limited phenotyping available. However, a substantial portion of that variability is likely reflective of experimental and/or measurement error. This variability can partially explain recent failures of replicability in neuroimaging [20], as well as the lack of clinically useful neuroimaging based biomarkers [27]. This work therefore motivates significant further efforts in measurement and/or analysis to mitigate “batch effects’’ in neuroimaging. Other disciplines with similar findings have resolved these issues by revolutionizing their measurement strategy (for example, genomics moved away from microarrays because sequencing can be significantly more reliable than microarray measurements [28])—though only after all efforts to remediate existing methods failed. For imaging, more comprehensively phenotyped individuals, and more coordinated data acquisition protocols, can be a first step towards studies generating sufficiently accurate and reliable inferences and clinical biomarkers.

**Table 1:**
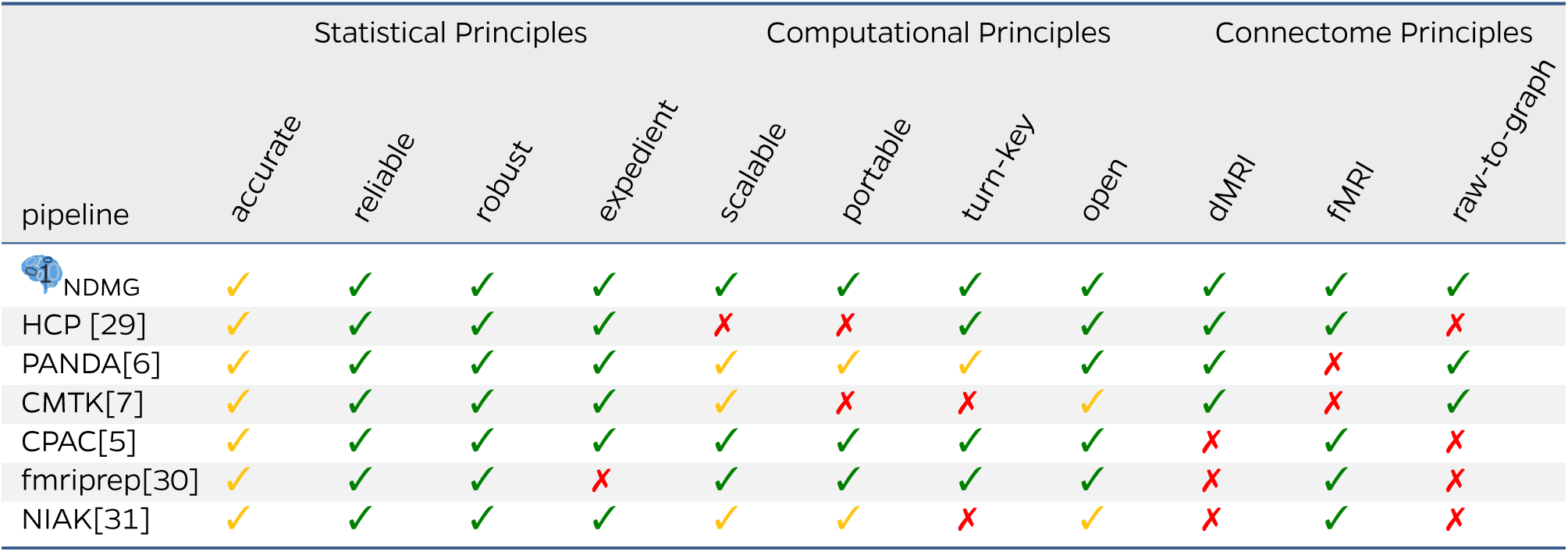
Comparing M3R Processing Pipelines. NDMG is designed with both algorithmic and implementation principles in mind. This table compares existing pipelines along these principles, demonstrating that for each, NDMG performs at least as well as the current state of the art. A ✔ is given for pipelines that satisfy the respective desiderata, a ✔ for pipelines that partially satisfy the respective desiderata, and a ✘ is given for pipelines that do not satisfy the respective desiderata. Appendix A provides more details.

## 2 Results

Table 1 depicts NDMG’s performance with regard to each of the below described principles for reproducible pipelines. For each principle, we outline a procedure for evaluating the degree to which a pipeline adheres to that principle, enabling the principle-based evaluation of disparate approaches.

### Statistical Principles

The statistical principles are designed to evaluate the empirical quality of the method on real data. Better quality suggests the pipeline can be trusted to be reliable for subsequent investigations.

*Accuracy* quantifies the distance between “the truth’ and an estimate—also known as “validity”—and is closely related to the statistical concepts of “unbiased” and “consistent”. Because the truth is unknown for these data, no pipeline can be meaningfully evaluated in terms of accuracy. Until the field develops the technological capability to obtain “ground truth” data, we rely on surrogate information. Specifically, NDMG incorporates the quality assessment (QA from CPAC [5] as well as other pipelines, and also adds several novel QA figures, with at least one figure per processing stage, yielding a total of 11 QA figures per diffusion scan and 32 per functional scan (see Appendix A for details). NDMG does not filter any data based on QA thresholds, as the choice of threshold remains both subjective and task dependent.

*Reliability* colloquially refers to methods that produce a similar result given a similar input, also called “stability” in the statistics literature [32]. To evaluate a method’s reliability, Wang et al. [33] developed a metric called “discriminability” that quantifies the fraction of measurements from the same individual that are closer to one another than they are to the measurement of any other individual (details below). NDMG’s discriminability over all scans was nearly 0.98 for dMRI data and over 0.88 for fMRI.

*Robustness* quantifies accuracy and reliability across a wide range of studies with different properties, including different experimental design, measurement devices, etc. We, therefore, ran NDMG on-dMRI studies and 18 fMRI studies using different hardware and acquisition parameters (see Table 2 for details). In some of these studies samples were not filtered to discard outliers or samples with poor signal-to-noise properties. Nonetheless, for each study, NDMG’s QA suggested accuracy, and each study achieved a score of discriminability > 0.8.

### Computational Principles

Adhering to the computational principles lowers the barrier for use. In practice, this means that both domain experts and researchers from other disciplines (such as machine learning and statistics), can more easily use the tools.

*Expediency* refers to the time it takes to run an approach on a typical sample. In practice, we suspect that users are more likely to adopt pipelines that run in about an hour per scan as compared to those that run overnight per scan, for example. The NDMG runtime is about 20 minutes for a functional scan, and 1.5 hours for a diffusion scan.

*Parallelized* refers to the ability of the method to parallelize the code across multiple computers. NDMG enables parallelization across scans—using the commercial cloud or a high-performance cluster, for example, enables NDMG to run many thousands of scans in only 1.5 hours.

*Portability—meaning* the ability to be run on different platforms, from laptop and cloud, with minimal installation and configuration energy—enables different analysts using different hardware resources to seamlessly use the code. We have tested NDMG on multiple hardware and operating system setups, including Windows, OSX, and Linux laptops, multi-core workstations, singularity clusters, and the Amazon cloud. Moreover, we have deployed NDMG on both openneuro [34] and CBRAIN [35], making it possible for anybody to run NDMG for free on their own or other’s computational resources.

*Turn-Key* methods do not require the user to specify parameters and settings for each stage of processing or for each new study. This feature reduces the time for researchers to get a pipeline running, and enables pooling data across multiple pipelines because the analysis is harmonized (conducted identically across studies). NDMG parameters have been tuned to yield accurate and reliable connectome estimates across nearly 30 different studies. Moreover, NDMG is fully compliant with Brain Imaging Data Structure (BIDS)—a recently proposed specification for organizing multi-scan, multi-individual, and multi-modality studies [36; 37].

*Open*, referring to both open source code and open access data derivatives processed using the code, enables anybody with Web-access to contribute to the scientific process. NDMG leverages open source packages with permissive licenses, and is released under the Apache 2.0 open source license. Our website, http://m2g.io contains links to download all of the data derivatives and quality assurance figures from each scan, and is the largest open database of connectomes and other data derivatives to our knowledge. By developing NDMG according to these statistical and computational design principles, and running it on many diverse studies, we can evaluate individuals, studies, and the collection of studies at an unprecedented scale. Below, we describe the nuts and bolts of the pipeline, followed by a set of NDMG -enabled scientific findings. Our hope is that NDMG and the data products derived from it will be useful for a wide variety of discoveries.

### Connectome Principles

We also consider a pair of principles that are specific for reproducible pipelines in connectomics.

*dMRI and fMRI* pipelines operate on diffusion or functional MRI data respectively.

*Raw-to-Graph* refers to the pipeline performing all steps of analysis required to acquire connectomes (graphs) given raw, unprocessed M3R scans with no user input. The NDMG-d pipeline was built to take raw dMRI and T1w images and produce a diffusion connectome, and the NDMG-f pipeline takes raw fMRI and T1w images and produces a functional connectome.

### 2.1 Individual-Level Analysis

In the individual-level analysis, each individual undergoes some number of sessions, during which multiple different modalities can be collected. The input to NDMG for a given individual is the collection of scans and some metadata for each scan, including a structural scan, and either, or both of, (1) a diffusion scan, including the diffusion parameters files, and (2) a functional scan, including the slice acquisition sequence. The individual-level of NDMG analysis leverages existing open source tools, including the fMRI Software Library (FSL) [38–40], Dipy [41], the MNI152 atlas [42], and a variety of parcellations defined in the MNI152 space [43–51] (see Appendix G for details about built-in parcellations included). All algorithms requiring hyper-parameter selection were initially set to the suggested parameters for each tool, and tuned to improve the accuracy, reliability, expediency, and robustness of the results. The output of each processing stage includes data derivatives and QA figures to enable individualized accuracy assessments. The QA figures at many stages include cross-sectional images at different depths in the three canonical planes (sagittal, coronal, and axial) of images or overlays. Example QA figures are provided in Appendix B and Appendix C.

**Figure 1:**
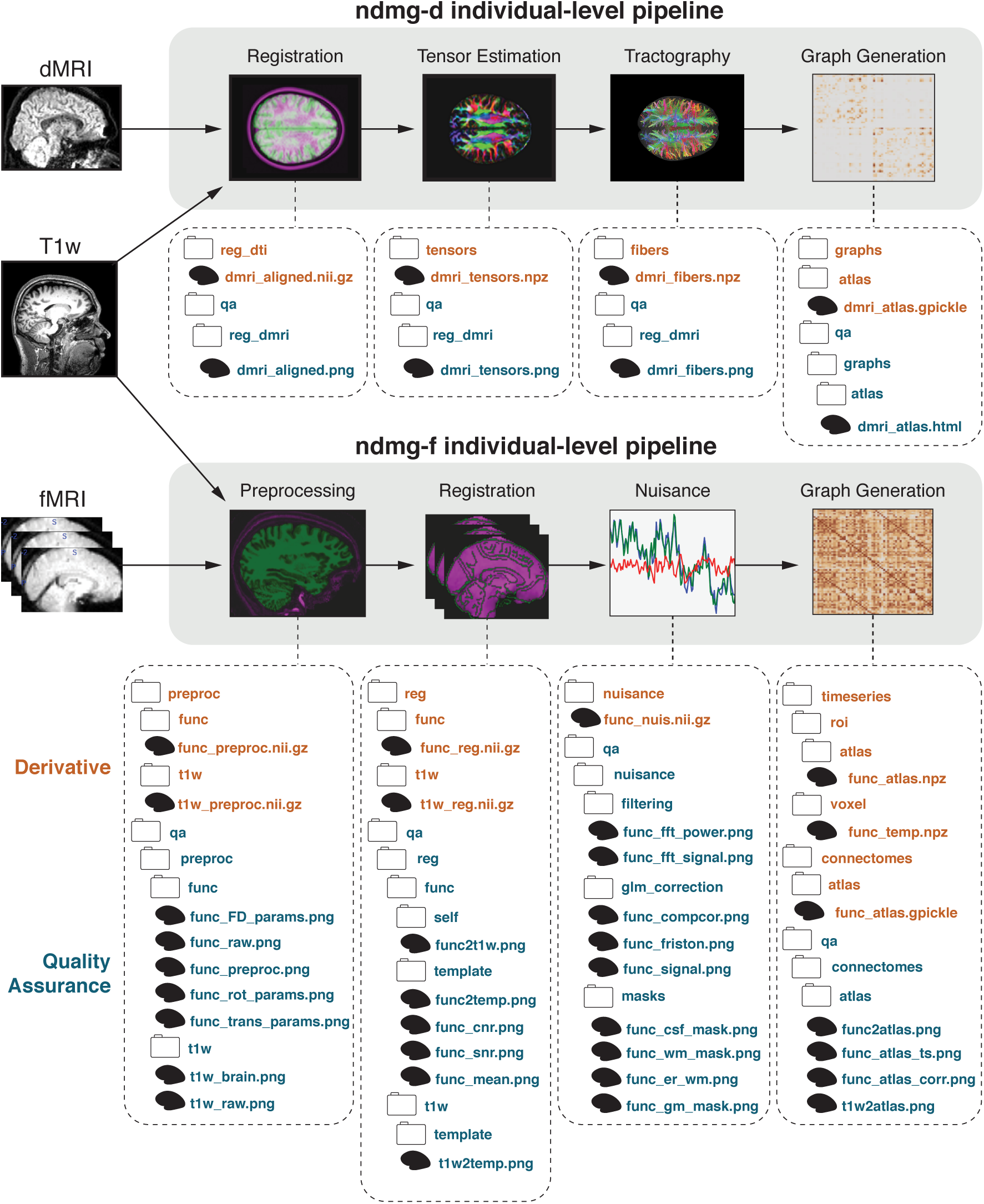
Individual Level Pipeline. The individual-level NDMG pipeline has two sub-pipelines: (1) NDMG-d transforms raw dMRI data into sparse structural connectomes, and (2) NDMG-f transforms raw fMRI data into dense functional connectomes. Each sub-pipeline consists of four key steps, and each step generates both data derivatives and quality assurance figures to enable both qualitative assessments and quantitative comparisons (see Appendix B and Appendix C for details).

#### Individual-Level Diffusion Analysis

The NDMG-d pipeline consists of four key components: (1) registration, (2) tensor estimation, (3) tractography, and (4) graph generation (see Figure 1 for an illustration, and Appendix B for further details). It was optimized on the Kirby21 dataset, and then applied to the remaning 10 datasets. Below, we provide a brief description of each step.

**Registration** uses FSL’s “standard” linear registration pipeline to register the structural and diffusion images to the MNI152 atlas [38–40; 42]. Nonlinear registration, though possibly more accurate [52; 53], was not sufficiently robust to include (it often failed on some studies without manual intervention).

**Tensor Estimation** uses DiPy [41] to obtain an estimated tensor in each voxel.

**Tractography** uses DiPy’s *EuDX* [54], a deterministic tractography algorithm closely related to *FACT* [55], to obtain a streamline from each voxel. Probabilistic tractography, while possibly more accurate, requires orders of magnitude more computational time. Visual QA of the generated fibers suggested that deterministic was sufficiently accurate for our purposes.

**Graph Generation** Graphs are formed by contracting fiber streamlines Appendix B.4 into sub-regions depending on spatial [56] proximity or neuro-anatomical [43–51] similarity. QA are performed as in the functional pipeline, described below.

Individual-level analysis in NDMG-d takes approximately 1.5 hours to complete using 1 CPU core and 12 GB of RAM at 1 mm^3^ resolution. The individual-level analysis was run on 11 studies, including 3,227 individuals and 4,347 scans. Each dataset generated connectomes across each of the parcellations in Appendix G, resulting in 104,328 functional brain-graphs.

#### Individual-Level Functional Analysis

The NDMG-f pipeline can be broken up into four key components: (1) preprocessing, (2) registration, (3) nuisance correction, and (4) graph generation (see Figure 1 for an illustration and Appendix C for further details). The NDMG-f pipeline was constructed starting with the optimal processing pipeline identified in Wang et. al [33] using CPAC [5]. Hyperparameters and further algorithm selection was optimized for reliability based on multiple measurement studies (including test-retest). Below, we provide a brief description of each step.

**Preprocessing** uses AFNI [57] for brain extraction, and FSL [58; 59] for slice-timing and motion correction.

**Registration** uses FSL [38; 60; 61] to perform a nonlinear boundary based registration of the fMRI into the MNI152 space [42]. The registration pipeline implemented is “standard” when working with functional data and FSL’s tools [5].

**Nuisance Correction** uses custom Python code to implement a general linear model incorporating regressors for quadratic detrending [62; 63], top five white-matter and cerebrospinal fluid Principal Components (CompCor) [64; 65], and the Friston 24-parameter regressors [66]. Low-frequency drift is then removed below 0.01 Hz [67], and the first 15 seconds of the fMRI sequence are discarded [68].

**Graph Generation** uses custom Python code to compute the average timeseries for all voxels within an ROI, then computes correlations between all pairs of ROIs. The functional connectome is then rank-transformed by replacing the magnitude of the correlation with its relative rank, from smallest to largest [33].

Individual-level analysis in NDMG-f takes approximately 20 minutes to complete using 1 CPU core and 3 GB of RAM at 2 mm^3^ resolution. The individual-level analysis was run on 714 individuals with 1,646 scans from 18 studies, generating connectomes across each of the 24 parcellations in NDMG-d, and resulting in 39,504 total brain-graphs.

#### Multi-Scale Multi-Connectome Analysis

For both diffusion and functional MRI, NDMG downsamples the voxel-wise graphs to obtain weighted graphs for many different parcellation schemes. This includes: (1) neuroanatomically delineated parcellations, such as the HarvardOxford cortical and sub-cortical atlases [46], the JHU [45], Talairach [47], Desikan [43], and AAL [44] atlases; (2) algorithmically delineated parcellations, such as Slab907 [48], Slab1068 [49], and CC200 [5]; and (3) 16 downsampled parcellations ranging from 70 to 72,783 nodes [56]. For each, the *multi-connectome* is defined by the set of nodes from a particular parcellation, and the set of (potentially weighted and/or directed) edges from each modality.

The QA for graph generation includes a heat map of the adjacency matrix, the number of non-zero edges, and several multivariate graph statistics (one statistic per vertex in the graph) including: betweenness centrality, clustering coefficient, hemisphere-separated degree sequence, edge weight, eigenvalues of the graph laplacian, and locality statistic-[56]. We developed the hemisphere-separated degree sequence to indicate the ipsilateral and contralateral degree for each vertex, which we found quite useful for QA. Appendix C.4 includes definitions and implementation details for each of the statistics. Supplementary Figure S10 shows, for a single individual, the graph summary statistics for the multi-connectome (including both functional and diffusion) across the set of atlases described above.

### 2.2 Group-Level Analysis

We ran NDMG on the 11 diffusion and 18 functional studies listed in Table 2. For each, NDMG group-level analysis computes and plots group-level graph summary and reliability statistics.

#### Group Level Implementation Strategy

Leveraging previous efforts developed in our “Science in the Cloud” [72] manuscript, multiple scans and studies are evaluated in parallel for participant-level analysis, and the derivatives are pooled for group-level analysis, much like typical map-reduce approaches (consistent with the BIDS app specification [37]). The parallel implementation uses the Amazon Web Services (AWS) cloud, in particular leveraging their storage (S3) and high performance computing (Batch) services. Data are stored in an S3 bucket enabling the NDMG cloud-API connectome estimation pipeline to process all scans on Amazon Batch in parallel. In practice, AWS limits the number of parallel threads users are allowed to launch (to prevent accidental spending). After connectome estimation is complete, the same cloud-API exposed by NDMG enables group-level analysis to be launched and parallelized across each parcellation for all scans within each study. We were able to compute diffusion connectomes at 24 scales for each of the publicly available 2,861 scans (totaling 68,664 connectomes) in under one day and $1,000. Had we processed each scan in parallel, cost would have remained the same but only taken 1.5 hours.

#### Group-Level Graph Summary Statistics

Each scan’s estimated connectome can be summarized by a set of graph statistics, as described above. For group-level analysis, we visualize each scan’s summary statistics overlaid on one another. For example, Figure 2 demonstrates that each diffusion graph from the BNU3 dataset using the Desikan atlas has relatively similar values for the statistics (we use the Desikan atlas for the remainder of the analyses unless otherwise specified). Moreover, it is clear from both the degree plot and the mean connectome plot that the dMRI connectomes from this study tend to have more connections within a hemisphere than across a hemisphere, as expected. For the fMRI connectomes, however, the homotopic connections—that is, connections from one region on a hemisphere to the same region on the other hemisphere—seem particularly strong.

#### Group-Level Discriminability

Group-level results from NDMG that include repeated measurements are quantitatively assessed using a statistic called discriminability [33]. The group’s sample discriminability estimates the probability that two observations within the same class are more similar to one another than to objects belonging to a different class:

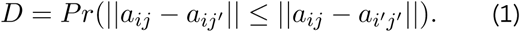

In the context of reliability in NDMG, each connectome, *a_ij_*, is compared to other connectomes belonging to the same individual, *a_ij′_*, and to all connectomes belonging to other individuals, *a_i′j′_*. A perfect discriminability score indicates that for all observations within the group, each connectome is more like connectomes from the same individual than others. Table 2 lists the discriminability score of each study with repeated measurements (five dMRI studies and sixteen fMRI). NDMG-d achieves a discriminability score of nearly 0.99 or greater on most studies (the lowest score was nearly 0.9). NDMG-f achieves a discriminability score around 0.9 on all studies. These high discriminability scores were achieved by optimizing both NDMG-d and NDMG-f on a subset of the data. Specifically, NDMG-d was optimized using only the Kirby21 study, to achieve a perfect discriminability score. Nonetheless, the other studies achieved comparably high discriminability scores, despite the fact that the Kirby21 used a Philips scanner, unlike any of the other studies. Moreover, Kirby21’s age distribution is substantially different from several of the other dMRI studies (see Table 2). Similarly, NDMG-f was optimized using only 13 of the 18 studies, and yet discriminability on the remaining studies remained equally high [33], even though they also exhibited large study demographic and acquisition protocol variability. Finally, NDMG was not optimized explicitly at all on multimodal data, nonetheless, for all datasets with multiple scans per subject with both dMRI and fMRI, using the multimodal connectomes improved (or did not decrease) discriminability relative to either modality on its own. That NDMG’s discriminability score is robust to data acquisition details and study demographics across modalities suggests that scientific results may also be robust to such sources of variability.

**Table 2:** NDMG pipeline robustness and reliability. We ran NDMG on over 20 different studies, including both fMRI and dMRI data, spanning multiple different scanners, acquisition parameters, and population demographics. Nonetheless, for both fMRI and dMRI data, across all datasets with multiple measurements, NDMG always achieved > 0.8 discriminability, and NDMG-d’s discriminability was typically > 0.98 on the dMRI data. MM computes discriminability using multi-modal data from both the dMRI and fMRI connectomes. *D*=discriminability. Scanners are G (GE), P (Phillips), or S (Siemens).

| Study | Scanner | Age±1 std | % Male | Indiv. | Ses. | dMRI | | fMRI | | MM | |
| --- | --- | --- | --- | --- | --- | --- | --- | --- | --- | --- | --- |
| | | | | | | #Scans | $D$ | #Scans | $D$ | #Scans | $D$ |
| HNU1 [21] | G | 24.4 ± 2.3 | 0.50 | 30 | 10 | 300 | 0.993 | 300 | 0.956 | 300 | .993 |
| BNU1 [21] | S | 23.0 ± 2.3 | 0.53 | 57 | 2 | 114 | 0.984 | 108 | 0.906 | 108 | .990 |
| SWU4 [21] | S | 20.0 ± 1.3 | 0.51 | 235 | 2 | 454 | 0.884 | 466 | 0.891 | 452 | .903 |
| BNU3 [21] | S | 22.5 ± 2.1 | 0.50 | 48 | 1 | 47 | NA | 47 | NA | 47 | NA |
| Kirby21 [69] | P | 31.8 ± 9.4 | 0.52 | 21 | 2 | 42 | 1.0 | - | - | - | - |
| NKI1 [21] | S | 34.4 ± 12.8 | 0.0 | 24 | 2 | 40 | 0.984 | - | - | - | - |
| Temp114 | S | 21.8 ± 3.0 | 0.58 | 114 | 1 | 114 | NA | - | - | - | - |
| NKI-ENH [70] | S | 42.5 ± 19.6 | 0.40 | 198 | 1 | 198 | NA | - | - | - | - |
| Temp255 | S | 22.1 ± 3.9 | 0.51 | 255 | 1 | 253 | NA | - | - | - | - |
| PING [71] | G/S/P | 11.7 ± 5.0 | 0.52 | 932 | 1-5 | 1486 | NA | - | - | - | - |
| MRN1313 | S | 27.6 ± 13.3 | 0.60 | 1313 | 1 | 1299 | NA | - | - | - | - |
| IPCAS6 [21] | S | 23.0 ± 2.0 | 0.5 | 2 | 9 | - | - | 18 | 0.994 | - | - |
| IPCAS8 [21] | S | 57.6 ± 3.6 | 0.46 | 13 | 2 | - | - | 26 | 0.958 | - | - |
| SWU1 [21] | S | 21.6 ± 1.7 | 0.3 | 20 | 3 | - | - | 60 | 0.935 | - | - |
| IPCAS5 [21] | S | 18.3 ± 0.5 | 1.0 | 22 | 2 | - | - | 44 | 0.819 | - | - |
| SWU3 [21] | S | 20.4 ± 1.6 | .35 | 23 | 2 | - | - | 46 | 0.986 | - | - |
| XHCUMS [21] | S | 52.0 ± 6.3 | 0.60 | 23 | 5 | - | - | 115 | 0.823 | - | - |
| UWM [21] | G | 25.0 ± 3.2 | 0.56 | 25 | 2 | - | - | 50 | 0.849 | - | - |
| NYU1 [21] | S | 29.4 ± 8.4 | 0.4 | 25 | 3 | - | - | 75 | 0.931 | - | - |
| SWU2 [21] | S | 21.0 ± 1.6 | 0.33 | 27 | 2 | - | - | 54 | 0.874 | - | - |
| IPCAS1 [21] | S | 20.9 ± 1.7 | 0.3 | 30 | 2 | - | - | 60 | 0.893 | - | - |
| IPCAS2 [21] | S | 13.4 ± 0.9 | 0.35 | 34 | 2 | - | - | 68 | 0.868 | - | - |
| IBATRT [21] | S | 28.0 ± 7.5 | 0.54 | 36 | 2 | - | - | 50 | 0.974 | - | - |
| MRN1 [21] | S | 15.1 ± 7.8 | 0.5 | 52 | 2 | - | - | 90 | 0.859 | - | - |
| BNU2 [21] | S | 20.9 ± .9 | 0.54 | 61 | 2 | - | - | 121 | 0.863 | - | - |
| Pooled dMRI |  |  |  | 3227 |  | 4347 | 0.979 | - | - | - | - |
| Pooled fMRI |  |  |  | 763 |  | - | - | 1798 | 0.881 | - | - |
| Pooled MM |  |  |  | 361 |  | - | - | - | - | 907 | 0.982 |

**Figure 2:**
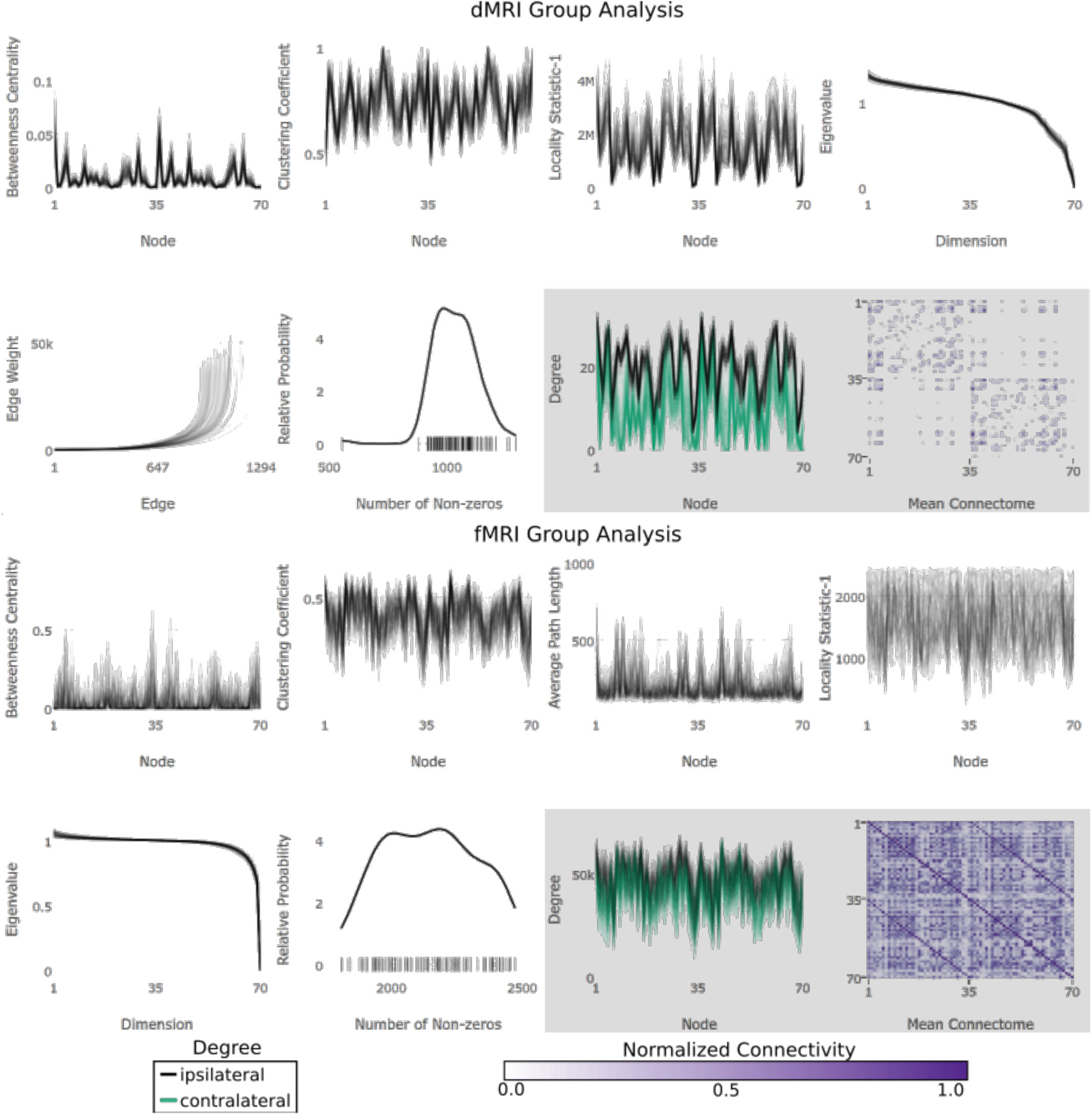
NDMG’s group level analysis computes and plots eight connectome-specific summary statistics per modality for each scan, providing immediate quality assurance for the entire study. In theory, any connectome could be an outlier for any of these statistics, so plotting all of them together is particularly useful (see Appendix D for details). The top panels show results for the BNU1 dMRI connectomes, the bottom panels show the results for the BNU1 fMRI connectomes.

### 2.3 Mega-Level Analysis

Many sources of variability contribute to the observed summary statistics, including individual- or population-level variation, and different types of measurement or analysis techniques. By virtue of harmonizing the analysis across individuals and studies, we are able assess the remaining degrees of variability due to measurement and demographic-specific effects. Although population-level effects are expected when comparing two different populations with different demographics, variability across measurements must be relatively small for inferences based on neuroimaging to be valid across these populations. Therefore, we conducted mega-analysis to address these remaining sources of variation.

#### Consistency Within and Across Studies

Figure 3 (top) shows the mean estimated connectome computed from each dataset for which we have both dMRI and fMRI data. We also calculate the mega-mean connectomes, that is, the mean derived by pooling all the studies (Figure 3, bottom). Several group-specific properties are readily apparent simply by visualizing the connectomes:

1. The dMRI connectomes have significantly stronger ipsilateral connections than contralateral connections, meaning connections are more prevalent within hemispheres, on average.
2. The fMRI connectomes have significantly stronger homotopic connections than heterotopic connections, meaning region A on the left hemisphere tends to correlation with region A on the right hemisphere, more than other regions, on average.

To formally test these initial assessments we developed statistical connectomics models and methods. Specifically, we developed a structured independent edge random graph model that generalizes the more commonly used stochastic block model (SBM) [73] and the independence edge random graph model [74]. In this new model, each edge is sampled independently, but not identically. Rather, there are *K* possible probabilities of an edge between a pair of vertices, and we have *a priori* knowledge of which edges are in which groups. Unlike the SBM model, in which each *vertex* is in a group, here each *edge* is in a group. We then developed test statistics that are consistent and efficient for this model. Specifically, with sufficiently large sample sizes, the power (the probability of correctly rejecting a false null hypothesis) of our test approaches unity for the model under consideration. Moreover, no other test can achieve higher power with fewer samples under this model (see Appendix E for details).

Using the above described approach, we first quantify the degree to which ipsilateral connections tend to be stronger than contralateral connections (Figure 4A). 100% of dMRI connectomes and 99.4% of fMRI connectomes, exhibit stronger ipsilateral than contralateral connections. The dMRI connectomes typically have a larger difference between ipsilateral and contralateral connections, as evidenced by the magnitudes of differences (Figure 4A.i). and the cumulative probability of a connectome having a difference that is statistically significantly at any level (Figure 4A.ii). We subsequently investigated whether for a given individual, the difference between ipsilateral and contralateral connections was stronger in the dMRI data than the fMRI data.

Out of 907 individuals with both dMRI and fMRI scans, 99.9% exhibit stronger ipsilateral versus contralateral connections in the dMRI scan as compared to their corresponding fMRI scan (Figure 4A.iii), with 99.5% significant at the 0.05 level, for example (Figure 4A.iv).

We applied the same strategy to test whether homotopic connections tend to be stronger than heterotopic connections (Figure 4B). In this case, the results are essentially opposite. Here, 100% of the fMRI and dMRI connectomes exhibit this property (Figure 4B.ii). Similarly, for nearly all individuals (99.9%) with both functional and diffusion scans, the relative strength of homotopic versus heterotopic connections was stronger in his or her fMRI data than the dMRI data (Figure 4B.iii), with 99.0% significant at the 0.05 level (Figure 4B.iv).

In other words, there is a marked consistency across *all* individuals in *all* studies (18 fMRI studies and 10 dMRI studies) for these most basic statistical properties of multi-modal connectomes. Notably, this result provides compelling evidence that certain connectomic discoveries that utilize only a single study, or even a single individual, without even addressing batch effects, can reasonably be expected to be repeatable across studies. Note, however, that repeatable does not mean correct; the true relative probabilities of ipsilateral, contralateral, homotopic, and heterotopic connections in humans remains an open question.

**Figure 3:**
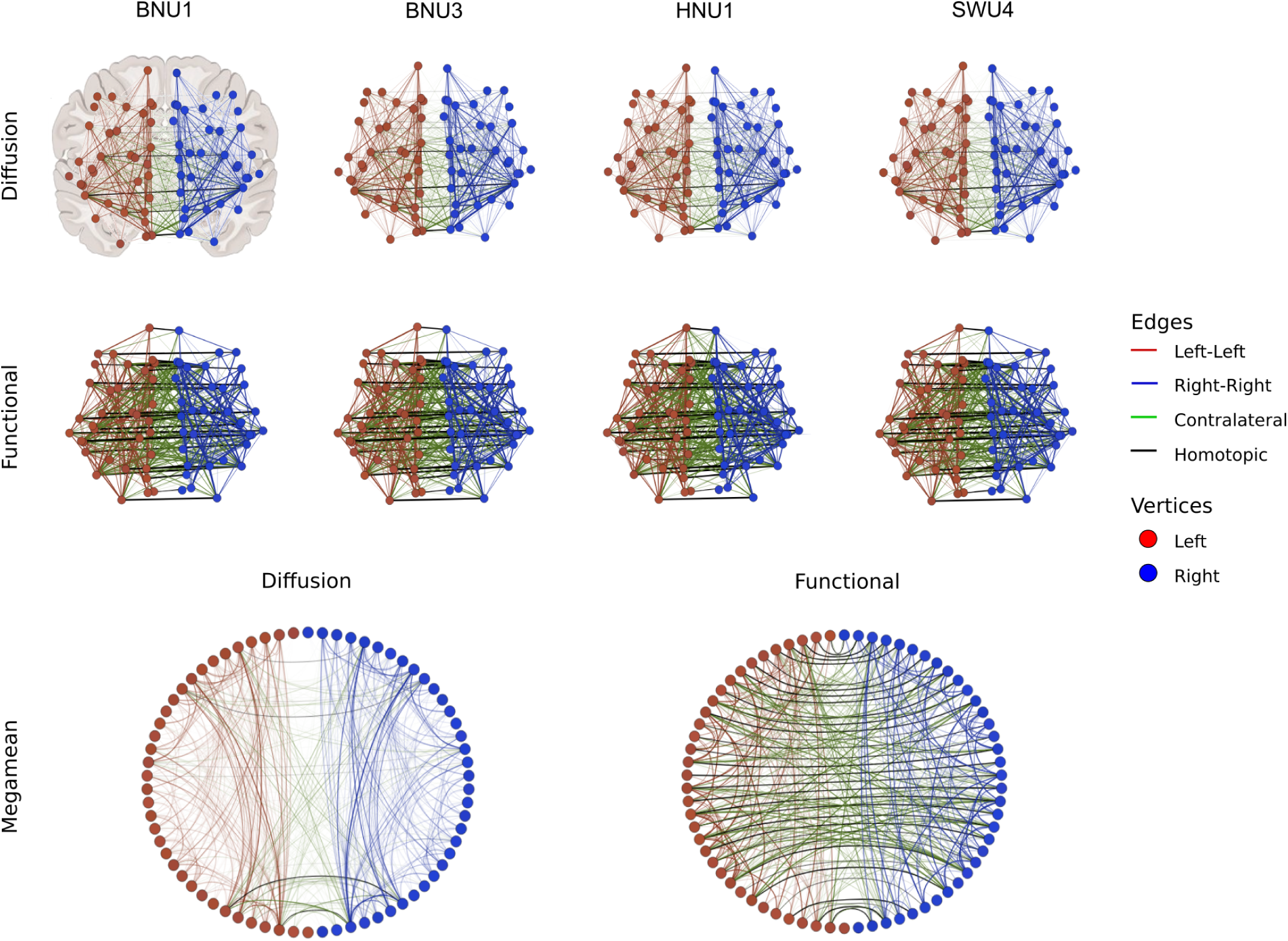
Multi-Study Mean Connectomes. Site-specific mean connectomes (top two rows) and mega-mean (bottom row), using the Desikan parcellation, using all the data for which we have both functional and diffusion studies (> 900 scans of each), with edges and vertices colored depending on hemisphere. In the top 2 rows, graphs are shown with vertex position determined by the coronally-projected center of mass for each region in the Desikan parcellation. The bottom shows radial plots organized by hemisphere. Same-modality connectomes appear qualitatively similar to one another across sites, but some differences across modalities are apparent. For example, homotopic and contralateral connections both seem stronger in functional than diffusion mean connectomes, both within site and after pooling all sites.

#### Significant Variability Within and Across Studies

While the above analysis demonstrates preservation of certain connectome properties both within and across studies, it is insufficient to determine the extent of “batch effects”—sources of variability including differences in participant recruitment, screening, and demographics as well as scanner, acquisition sequence, and operator. Typically, investigators prefer that their results are robust to these additional sources of variance. If the batch effects are larger than the signal of interest (for example, whether a particular individual is suffering from a particular psychiatric disorder), then inferences based on individual studies are prone to be unreliable.

**Figure 4:**
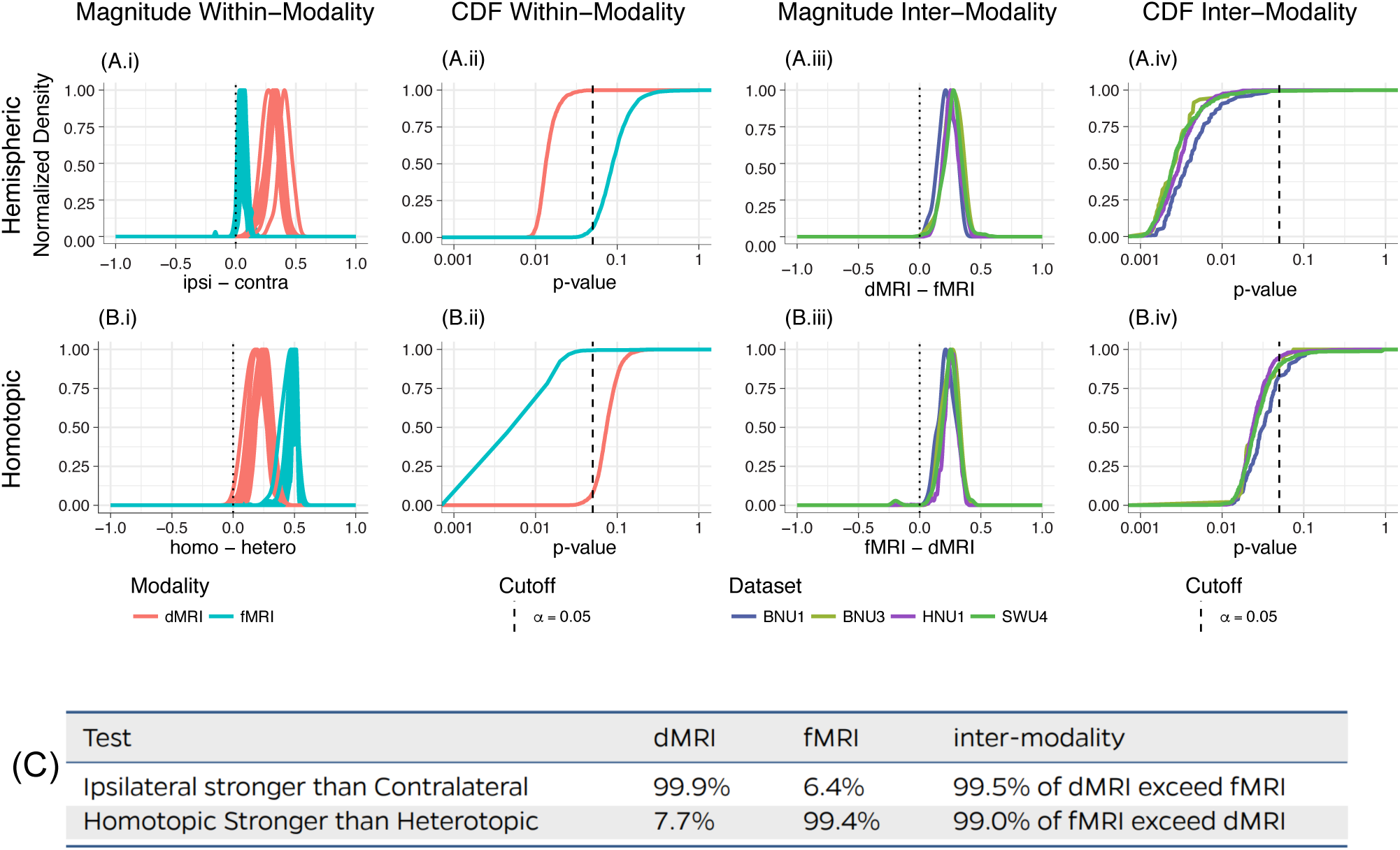
Structured Independent Edge Model (SIEM) establishes multi-connectome properties are preserved both within and across studies. **(A.i)** The magnitude of the difference indicating whether ipsilateral (within hemisphere) connections differ from contralateral (across hemisphere) connections. The dMRI connectomes appear to have a greater connectivity difference than fMRI connectomes. **(A.ii)** The distribution of p-values indicating whether ipsilateral connections are significantly stronger than contralateral connections, on average. Essentially every dMRI connectome yields significantly stronger ipsi- than contralateral connectivity, whereas essentially no fMRI connectome does. **(A.iii)** For four studies with both dMRI and fMRI, one can compare connection strengths across modalities for a particular individual. The ipsi-versus-contra discrepancy in the dMRI exceeds that of the fMRI. **(A.iv)** In essentially all sessions, dMRI has a significantly stronger within-versus-across discrepancy of connection strength than the corresponding fMRI connectome in most of same-individual pairs. **(B.i)** Same analysis comparing homo- to heterotopic connections, indicating that the fMRI connectomes appear to have a greater connectivity difference than the dMRI connectomes. **(B.ii)** fMRI exhibits significantly stronger homotopic connections, whereas dMRI does not. **(B.iii)** Same approach as **(A.iii)**, the homo-versus-hetero discrepency in the fMRI exceeds that of the dMRI. **(B.iv)** fMRI has a significantly stronger homo-versus-hetero discrepancy of connection strength than the corresponding dMRI connectome. **(C)** Table showing fraction of individuals exhibiting significance effects of each analysis described above, demonstrating consistency across individuals and studies.

Figure 5 evaluates the variability of individual scans both within and across studies for dMRI (left) and fMRI (right). For simplicity, we focus on a single parameter for each modality: dMRI data is evaluated in terms of its average within hemisphere connectivity and fMRI is evaluated in terms of its average homotopic connectivity. Prior to analysis each graph is “rank normalized”, meaning that its edge weights are converted into the numbers 1, 2,…, *N*, where *N* is the total number of potential edges, and where the smallest value is assigned a 1, the next smallest is assigned a 2, etc. By virtue of this normalization, each network has the same mean and variance, therefore mitigating certain kinds of batch effects. Moreover, by partitioning edges into only two groups, there is only a single degree of freedom: the average connection strength from one group determines that of the other, and vice versa (see Appendix F for details). The fact that the most salient features when visualizing these connectomes effectively yields a one-dimensional characterization of this property justifies the importance of studying the batch effects of this parameter. Dramatic variability in this parameter suggests that more “fine” parameters, such as the connection strength between individual regions of interest, will typically necessarily have even greater variability.

**Figure 5:**
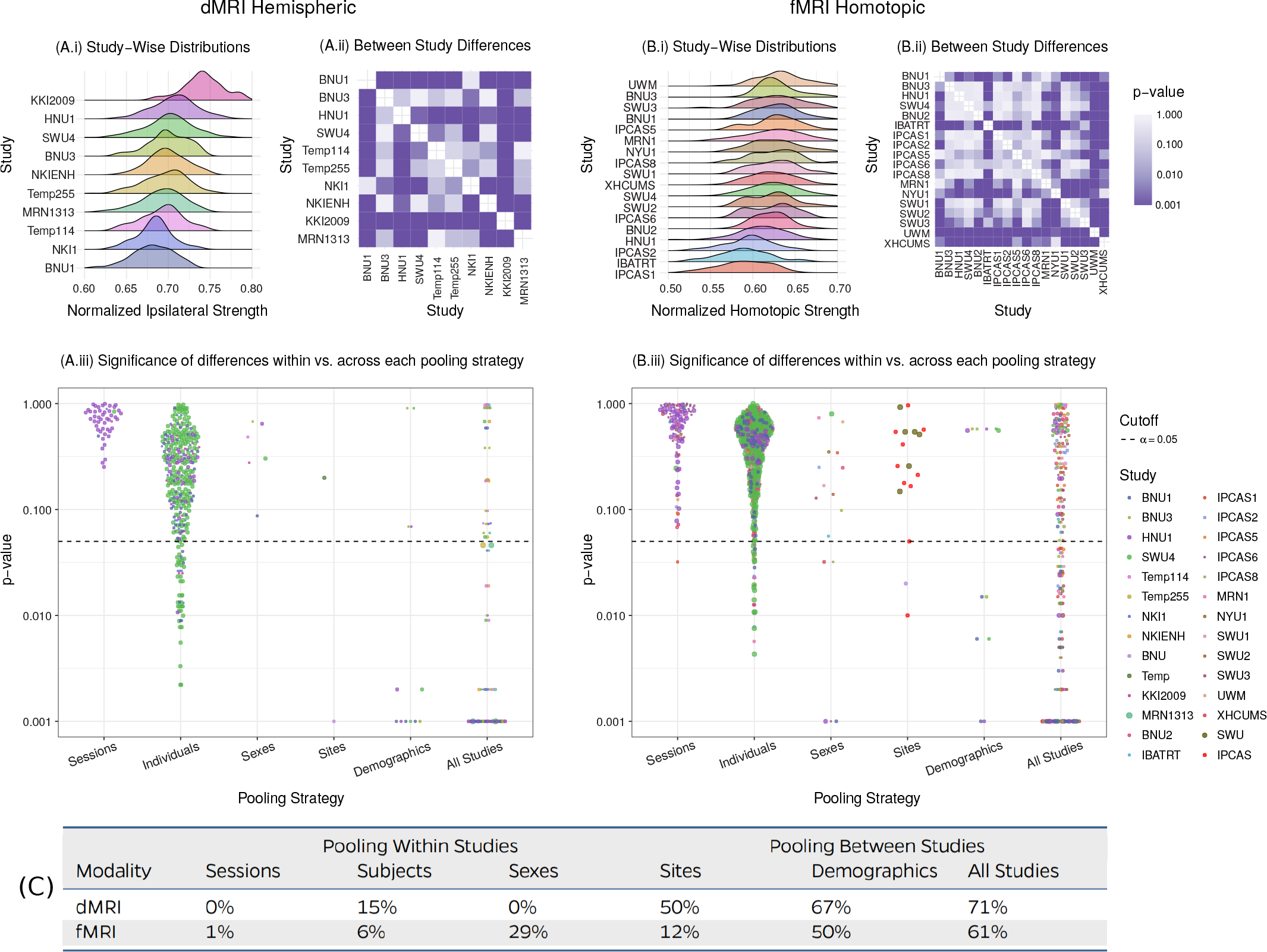
The prevalence of connectome batch effects are investigated within and across studies using different delineations of the edges into communities for dMRI and fMRI. **(A)** dMRI connectomes delineate edges into within and across hemisphere. **(B)** fMRI connectomes delineate edges into homotopic and heterotopic edges. **(A.i)** and **(B.i)** Density estimates of distribution of average ipsilateral or homotopic connection strength within studies. Both within and across study variance is distressingly high. **(A.ii)** and **(B.ii)** The pairwise significance of the differences between studies demonstrates that many pairs of studies exhibit significant batch effects. **(A.iii)** and **(B.iii)** Several common pooling strategies are investigated, including pooling across *sessions* within a study, pooling across *subjects* within a study, pooling across sex within a study, pooling across studies from a scanning *site*, pooling across studies with a similar basic *demographics*, and pooling across *all studies*. In many cases, even when controlling for these factors, significant batch effects remain. **(C)** The fraction of samples showing significant batch effect at *α* = 0.05. Pooling across sessions, subjects, and sexes, a small fraction of the samples show significant batch effect in both the dMRI and fMRI connectomes. Pooling across demographics and across all studies shows large batch effects. These results indicate that multiple (typically uncontrolled) variables considerably impact connectome inferences, implying that further efforts to mitigate these effects will be required to obtain sufficiently reliable estimates.

Figure 5(A.i) and (B.i) shows fairly dramatic variability both within and across studies, suggesting the presence of significant batch effects in both cases. Figure 5(A.ii) and (B.ii) evaluates the statistical significance of batch effects across studies, demonstrating that many studies, though not all, do exhibit severe batch effects. In this case, batch effect was quantified by determining whether the distribution of connection strengths for one study was significantly different from those of another study.

To further understand the source of this variability, we conducted an extensive within and between study analysis of individuals, including the following six different cases (Figure 5(A.iii) and (B.iii)):

1. Across *sessions* for a given individual within studies: If session 1 scans tended to be different from session 2 scans within a study, then the “scan order” itself can meaningfully impact inferences. While dMRI sessions never resulted in significant scan order effects, 9% of fMRI studies were significantly impacted by this effect.
2. Across *individuals* within studies: This analysis quantifies the fraction of individuals in a given study whose scans are less similar to one another than they are to any other scan in the dataset. 6% and 15% of individuals within studies (for fMRI and dMRI, respectively) have significantly different connection strengths.
3. Across *sexes* within studies: Connection strengths are the same at this coarse level across sexes for each dMRI study, whereas 30% of fMRI studies demonstrated significant differences across sexes.
4. Across studies within *sites:* Two dMRI and three fMRI sites provided data from multiple studies; thus, while the location and scanner were the same across these studies, certain variability in demographic and acquisition details persisted. Both dMRI and fMRI demonstrated a mix: sometimes these differences were significant, but not always.
5. Across studies within *demographic:* four dMRI and four fMRI studies controlled basic demographic variables, including only Chinese people, about 50% female, and all college age. Even preserving these demographics was typically insufficient to mitigate batch effects, with 2/3 and 1/2 of comparisons yielding significant batch effects for dMRI and fMRI, respectively.
6. Across *all* studies: 10 dMRI and 18 fMRI studies were compared, ignoring acquisition and demographic detail. Over 60% of pairwise comparisons for both dMRI and fMRI demonstrated significant differences, often the maximal significance level given our permutation test.

Taken together, the above results indicate the existence of many difference sources of variability, even upon harmonizing analyses (Figure 5(C)). Such variability suggests that the reproducibility of both dMRI and fMRI can benefit by further understanding, quantifying, and mitigating these sources of variability.

## 3 Discussion

The goal of any scientific investigation is not to characterize the observed sample data’s variance, but rather, to make inferences about the general population on the basis of those data. Variability of sample demographics, data acquisition details (for example, number of repetitions for fMRI, number of directions for dMRI), analysis methods, measurement error, questionable research practices, or statistical errors can each contribute to limitations in generalizing to populations in psychology [20] and neuroimaging [75]. Our principles for pipeline development enabled a rigorous high-throughput analysis of multi-study, multi-site, and multi-connectome data to identify and quantify different sources of variability.

While perhaps seemingly at odds, the results from Figures 4 and 5 are complementary. Specifically, Figure 4 demonstrates that essentially all connectomes share a particular property: stronger connections in one set of edges than another. On the other hand, Figure 5 demonstrates that although some sets of edges tend to be larger than others, the magnitude of the differences is highly variable both within and across studies. That the *magnitude* of certain parameters can differ while the *sign* of those parameters can be constant parallels recent suggestions from the statistics literature to move away from “Type I” and “Type II” errors, to Type M (magnitude) and Type S (sign) errors [76]. Moreover, it suggests a strategy to understand and address batch effects: reporting the *precision* for which effects are preserved or variable. For example, in this study, when using the coarsest possible precision (larger versus smaller as in Figure 4), no batch effects arise, whereas when using an extremely fine precision (the difference in magnitude as in Figure 5), batch effects are pervasive. The natural question then becomes: at which precision, for a given parameter and source of variability, do the studies still agree? Such analyses could then be incorporated into downstream analyses to preserve results across studies.

The design criteria for NDMG required certain trade-offs, including robustness under variability versus optimality under idealized data. Nonetheless, NDMG, could be improved along several dimensions. First, recent advances in registration [53] and tractography [77] could be incorporated. When implementations for these algorithms achieve suitable expediency and robustness, it will be natural to assess them. Second, recently several more sophisticated batch effect strategies have been successfully employed in dMRI data [78]. Such strategies could possibly help here as well, especially if they are modified appropriately to work on binary graphs [79], Third, there is some evidence that machine learning approaches to mitigating batch effects can be effective as well, but so far only in fMRI data and only using four, rather than 18 studies [80]. Fourth, pre-processing strategies have been employed to improve multi-site reliability [81], so implementing methods such as these within NDMG could possibly mitigate some batch effects, at the risk of reducing accuracy and/or reliability [82].

It may be that analysis methods on their own are insufficient to mitigate replicability issues, and that further improving data acquisition and/or data acquisition harmonization may be required. Indeed, a recent study by Noble et al. [83] found relatively few batch effects in fMRI data, although it employed only two datasets with enhanced and harmonized data acquisition protocols.

Because the methods developed herein are open source and easy to use, and the data are open access, this work enables further studies to assess measurement errors as well as variability of sample demographics and experimental protocols. For example, data could be sub-sampled to only include scans that pass stringent quality assurance standards, or have a sufficiently long duration to support discriminable connectomes [**?**]. Alternately, this analysis could be repeated on data that is “perfectly” harmonized. In general, further work developing and applying experimental and theoretical tools to parse the relative impact of various sources of batch effects, as well as batch effect mitigation strategies, will be crucial for neuroimaging to achieve its greatest potential scientific and clinical value.

## Acknowledgements

The authors from JHU are grateful for the support by the XDATA program of the Defense Advanced Research Projects Agency (DARPA) administered through Air Force Research Laboratory contract FA8750-12-2-0303; DARPA SIMPLEX program through SPAWAR contract N66001-15-C-4041; DARPA GRAPHS contract N66001-14-1-4028; National Science Foundation grant 1649880, and the Kavli Foundation for their support. We are grateful to Eric Walker^1,5^ and Tanay Agarwal^1,5,14^ who helped with preliminary QA figures for NDMG-f. Dr. Xi-Nian Zuo received funding support in China from the National Basic Research (973) Program (2015CB351702), the National R&D Infrastructure and Facility Development Program “Fundamental Science Data Sharing Platform” (DKA2017-12-02-21), the Natural Science Foundation of China (81471740, 81220108014) and Beijing Municipal Science and Tech Commission (Z161100002616023, Z161100000216152). Dr. Calhoun received funding from the NIH (P20GM103472 and R01EB020407) and the NSF (grant 1539067).

## CoRR Members

Jeffrey S. Anderson, Pierre Bellec, Rasmus M. Birn, Bharat B. Biswal, Janusch Blautzik, John C.S. Breitner, Randy L. Buckner, F. Xavier Castellanos, Antao Chen, Bing Chen, Jiangtao Chen, Xu Chen, Stanley J. Colcombe, William Courtney, Adriana Di Martino, Hao-Ming Dong, Xiaolan Fu, Qiyong Gong, Krzysztof J. Gorgolewski, Ying Han, Ye He, Yong He, Erica Ho, Avram Holmes, Xiao-Hui Hou, Jeremy Huckins, Tianzi Jiang, Yi Jiang, William Kelley, Clare Kelly, Margaret King, Stephen M. LaConte, Janet E. Lainhart, Xu Lei, Hui-Jie Li, Kaiming Li, Kuncheng Li, Qixiang Lin, Dongqiang Liu, Jia Liu, Xun Liu, Guangming Lu, Jie Lu, Beatriz Luna, Jing Luo, Daniel Lurie, Ying Mao, Andrew R. Mayer, Thomas Meindl, Mary E. Meyerand, Weizhi Nan, Jared A. Nielsen, David O’Connor, David Paulsen, Vivek Prabhakaran, Zhigang Qi, Jiang Qiu, Chunhong Shao, Zarrar Shehzad, Weijun Tang, Arno Villringer, Huiling Wang, Kai Wang, Dongtao Wei, Gao-Xia Wei, Xu-Chu Weng, Xuehai Wu, Ting Xu, Ning Yang, Zhi Yang, Yu-Feng Zang, Lei Zhang, Qinglin Zhang, Zhe Zhang, Zhiqiang Zhang, Ke Zhao, Zonglei Zhen, Yuan Zhou, Xing-Ting Zhu.

## Appendix A Pipeline Comparison Technology Evaluation

The scoring criteria for the principles defined in the main text are defined as follows:

**Accurate** ✔: pipeline results compared with ground truth to assess accuracy, ✔: quality control figures and metrics are produced.

**Reliable** ✔: reliability has been assessed, using either discriminability or intra-class correlation, by running the pipeline on at least one study, ✔: no published results demonstrating reliability

**Robust** ✔: pipeline has been run successfully on multiple disparate studies, ✔: no published results on multiple studies.

**Expedient** ✔: pipeline runs on a single individual ≤ 2 hour per scan, ✘: pipeline requires ≥ 2 hour per scan.

**Parallelized** ✔: can run in parallel locally, and can scale to cloud infrastructure (AWS EC2 or AWS Batch). ✔: can run in parallel locally or scale to cloud infrastructure. ✘ : can neither run in parallel locally nor scale to cloud infrastructure.

**Portable** ✔: can run on, and is deployed on, multiple different platforms, ✔: can run on multiple platforms, but is not deployed on any publicly available resources, ✘ : is platform specific.

**Turn-Key** ✔: runs without requiring any tuning parameters or configuration files, ✔: given a configuration file, runs without requiring any further tuning, ✘ : requires tuning for each run.

**Open** ✔: both source code and data derivatives are open, ✔: only source code is available, ✘ : neither source code nor data derivatives are publicly available.

**dMRI&fMRI** ✔ operates on dMRI data (left), operates on fMRI (right), ✘ on either side indicates the opposite.

**Raw-to-Graph** ✔: outputs estimated networks, ✘ : does not.

## Appendix B Diffusion Pipeline

Here we take a deep-dive into each of the modules of the NDMG-d pipeline. We will explain algorithm and parameter choices that were implemented at each step and the justification for why they were used over alternatives. All QA/QAX figures shown are from participant 0025864 from the BNU1 [21] study.

**Table 3:** NDMG-d IO Breakdown. Below, we look at the inputs, outputs, and QA figures produced by NDMG-d.

| Step | Inputs | Outputs | QA figures |
| --- | --- | --- | --- |
| Registration | raw dMRI, raw T1w, template | aligned dMRI | Figure S2 |
| Tensor Estimation | aligned dMRI | tensor field | Figure S3 |
| Tractography | tensor field | fiber tracts | Figure S4 |
| Graph Generation | fiber tracts, parcellations (Appendix G) | connectome |  |

### Appendix B.1 Registration

Registration in NDMG leverages FSL and the Nilearn Python package. NDMG uses linear registrations because non-linear methods had higher variability across studies and increased the time requirements of the pipeline dramatically (not shown).

The first step in the registration module is eddy-current correction and dMRI self-alignment to the volume-stack’s BO volume (Figure S1). NDMG uses FSL’s `eddy_correct`, rather than the newer `eddy` function, because `eddy` requires substantially longer to run or relies on GPU acceleration, which would reduce the accessibility of NDMG. Once the dMRI data is self-aligned, it is aligned to the same-individual T1w image through FSL’s `epi_reg` mini-pipeline. This tool performs a linear alignment between each image in the dMRI volume-stack and the T1w volume. The T1w volume is then aligned to the MNI152 template using linear registration computed by FSL’s `flirt`. This alignment is computed using the 1 millimeter (mm) MNI152 atlas, to enable higher freedom in terms of the parcellations that may be used, such as near-voxelwise parcellations that have been generated at 1 mm. FSL’s non-linear registration, `fnirt`, is not used in NDMG as the performance was found to vary significantly based on the collection protocol of the T1w images, often resulting in either slightly improved or significantly deteriorated performance.

The transform mapping the T1w volume to the template is then applied to the dMRI image stack, resulting in the dMRI image being aligned to the MNI152 template in *stereotaxic*-coordinate space. However, while `flirt` aligns the images in stereotaxic space, it does not guarantee an overlap of the data in voxelspace. Using Nilearn’s `resample`, NDMG ensures that images are aligned in both voxel- and stereotaxic-coordinates so that all analyses can be performed equivalently either with or without considering the image affine-transforms mapping the data matrix to the real-world coordinates.

**Figure S1:**
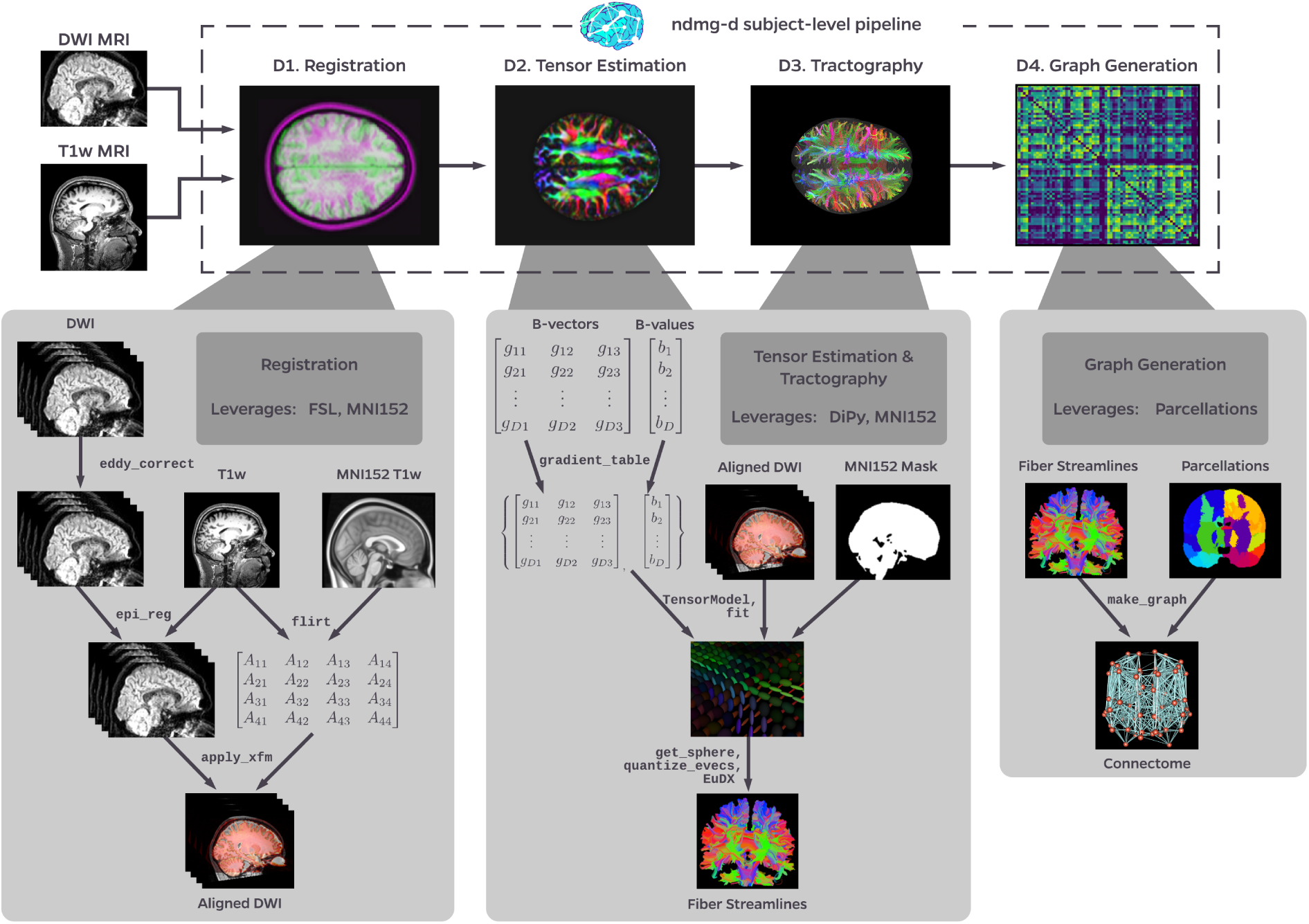
NDMG-d detailed pipeline. The NDMG-d pipeline consists of 4 main steps: Registration **(D1)**, Tensor Estimation **(D2)**, Tractography **(D3)**, and Graph Generation **(D4)**. Each of these sections leverages pubicly available tools and data to robustly produce the desired derivative of each step. Alongside derivative production, NDMG produces QA figures at each stage, as can be seen in **D1-4**, that enable qualitative evaluation of the pipeline’s performance.

**Figure S2:**
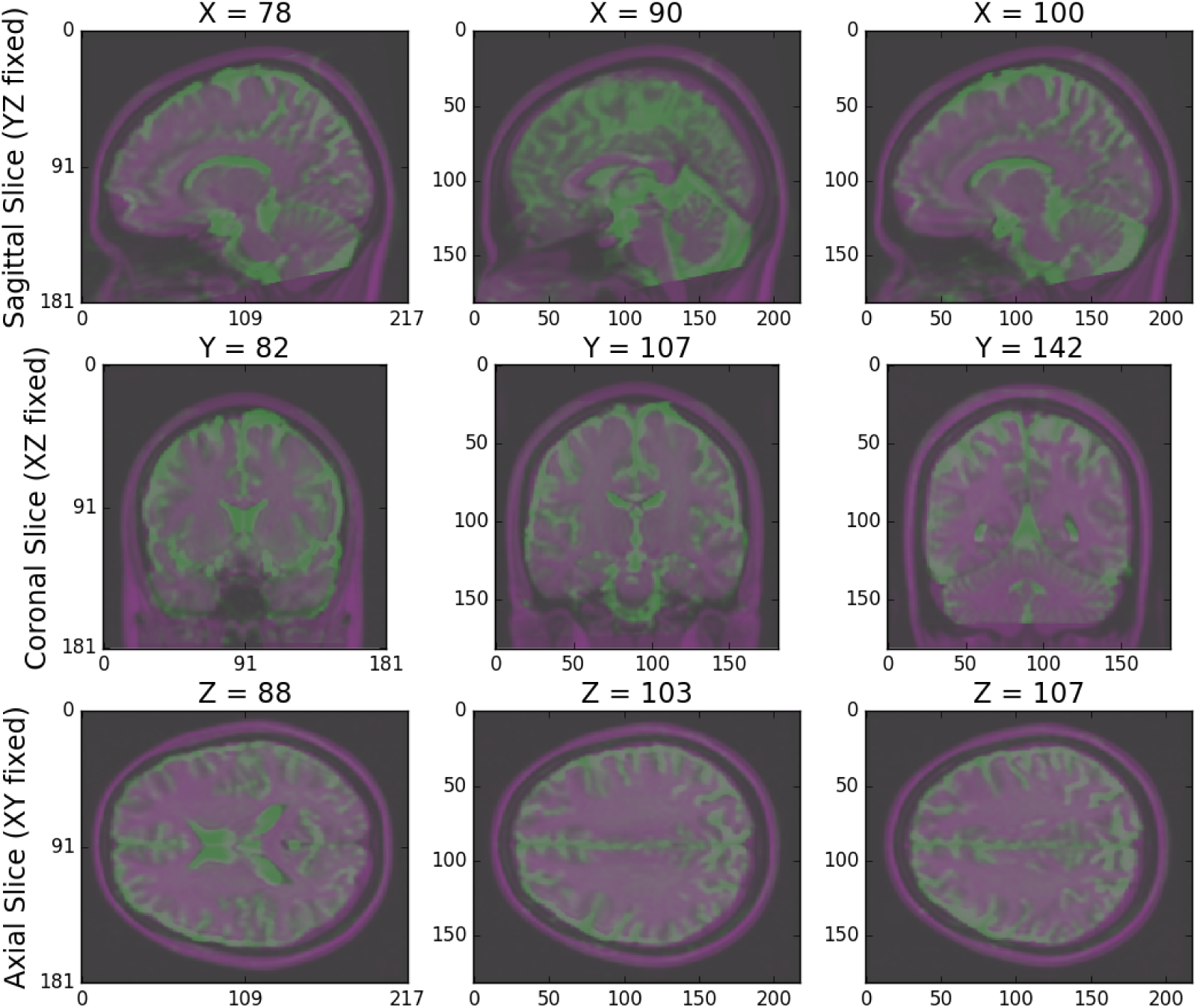
NDMG-d Registration QA. NDMG-d produces registration QA showing the zeroth slice of the dMRI sequence in green overlaid on the template brain in purple.

**Figure S3:**
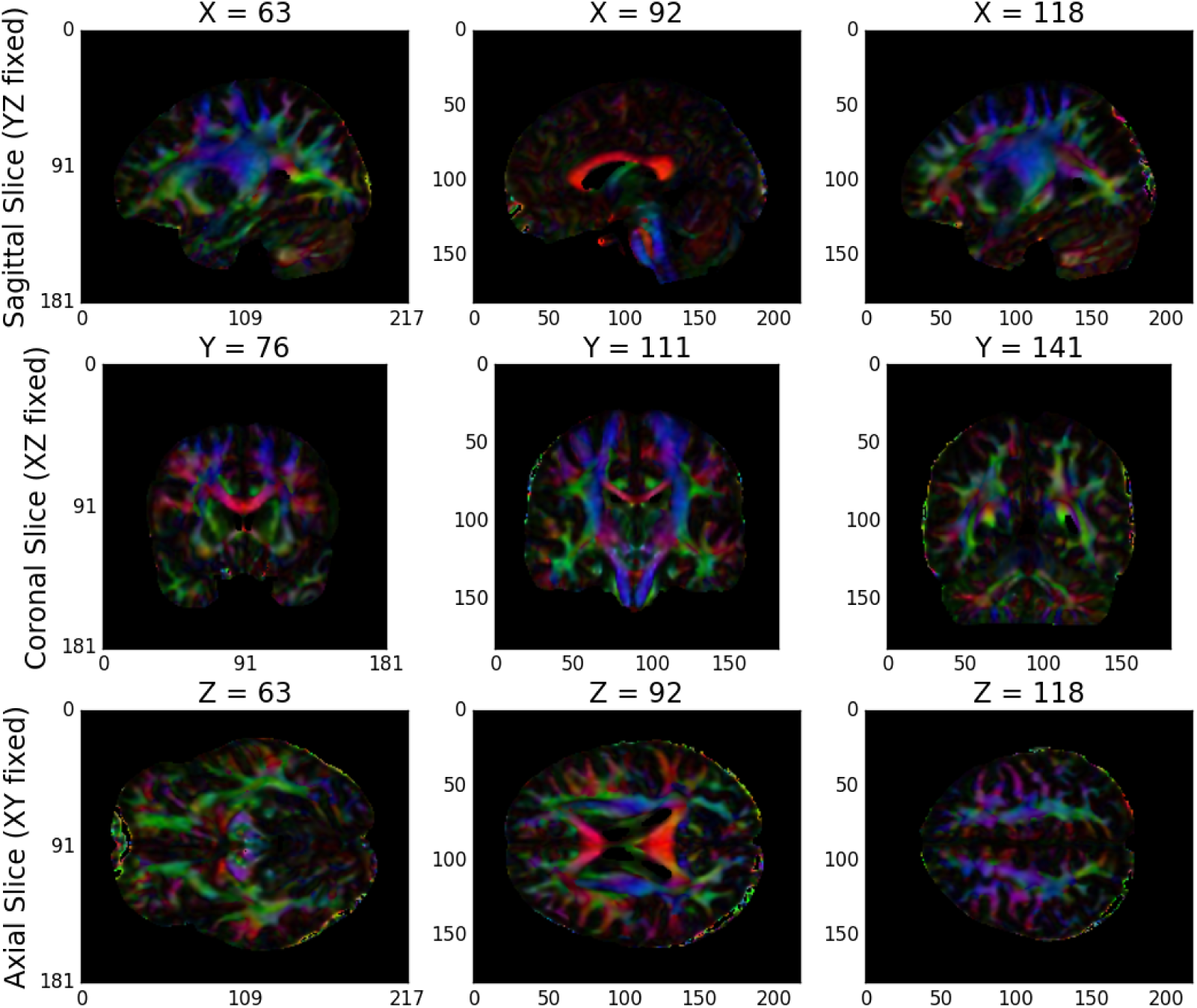
NDMG-d Tensor Estimation QA. NDMG-d produces tensor QA showing the voxelwise determininstic tensor model fit to the aligned dMRI sequence.

Finally, NDMG produces a QA plot showing three slices of the first B0 volume of the aligned dMRI image overlaid on the MNI152 template in the three principle coordinate planes (Figure S2).

### Appendix B.2 Tensor Estimation

Once the dMRI volumes have been aligned to the template, NDMG begins diffusion-specific processing on the data. All diffusion processing in NDMG is performed using the Dipy Python package [41]. The diffusion processing in NDMG is performed after alignment to ease cross-connectome comparisons.

While high-dimensional diffusion models, such as orientation distribution functions (ODFs) or q-ball, enable reconstruction of crossing fibers and complex fiber trajectories, these methods are designed for images with a large number of diffusion volumes/directions for a given image [84; 85]. Because NDMG is designed to be robust across a wide range of dMRI studies, including diffusion tensor imaging, NDMG uses a lower-dimensional tensor model. The model, described in detail on Dipy’s website (http://nipy.org/dipy/examples_bcouilt/reconst_dti.html), computes a 6-component tensor for each voxel in the image. This reduces the dMRI image stack to a single 6-dimensional image that can be used for tractography. NDMG generates a QA plot showing slices of the FA map derived from the tensors in nine panels, as above (Figure S3).

**Figure S4:**
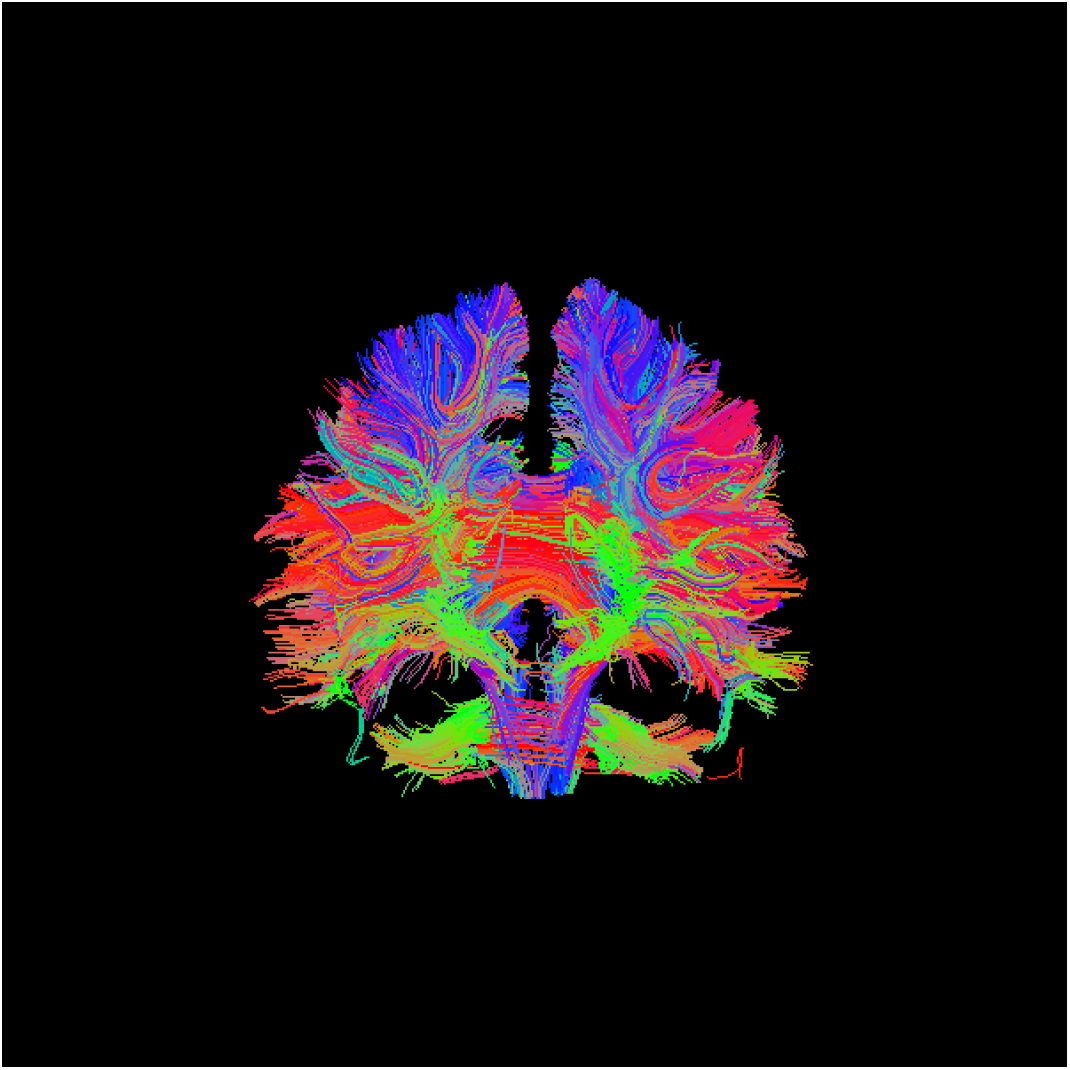
NDMG-d Tractography QA. NDMG-d produces tensor QA showing the fiber tracts in the brain.

### Appendix B.3 Tractography

NDMG uses DiPy’s deterministic tractography algorithm, `EuDX` [54]. Integration of tensor estimation and tractography methods is minimally complex with this tractography method, as it has been designed to operate on the tensors produced by Dipy in the previous step. Probabilistic tractography would be significantly more computationally expensive, and it remains unclear how well it would perform on data with a small number of diffusion directions. The QA figure for tractography shows a subset of the resolved streamlines in an axial projection of the brain mask with the fibers contained therein (Figure S4). This QA figure allows the user to verify, for example, that streamlines are following expected patterns within the brain and do not leave the boundary of the mask.

### Appendix B.4 Graph Estimation

NDMG uses the fiber streamlines to generate connectomes across multiple parcellations. Graphs are formed by contracting voxels into graph vertices depending on spatial [56], anatomical [], or functional [] similarity. Given a parcellation with vertices *V* and a corresponding mapping *P*(*v_i_*) indicating the voxels within a region *i*, we contract our fiber streamlines as follows. To form a graph *G*(*V, E,w*), we know that 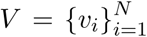 is the set of all unique vertices in our parcellation. *E* = *V* × *V* is a collection of possible edges between different vertex regions, *w*: *V* × *V* → ℤ_+_ is a weighting function for each edge in the graph. Here, *w*(*v_i_, v_j_*) = 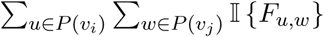 where *F_u,w_* is true if a fiber tract exists between voxels *u* and *w*, and false if there is no fiber tract between voxels *u* and *w*.

The connectomes generated are graph objects, with nodes in the graph representing regions of interest (ROIs), and edges representing connectivity via fibers. An undirected edge is added to the graph for each pair of ROIs a given streamline passes through. Edges are undirected because dMRI data lacks direction information. Edge weight is the number of streamlines which pass through a given pair of regions. NDMG uses 24 parcellations, including all standard public dMRI parcellations known by the authors. Users may run NDMG using any additional parcellation defined in MNI152 space simply by providing access to it on the command-line. To package an additional parcellation with NDMG, please contact the maintainers. The QA for graph generation depicts a number of graph statistics for each of the parcellation schemes. We typically generate this figure at the population level, as depicted in Figure 2. The description of all the graph statistics we used is provided in Appendix D.

**Table 4:** NDMG-f IO Breakdown. Below, we look at the inputs, outputs, and QA figures produced by NDMG-f.

| Step | Inputs | Outputs | QA figures |
| --- | --- | --- | --- |
| Preprocessing | raw fMRI, raw T1w | preprocessed fMRI, preprocessed T1w | Figure S6 |
| Registration | preprocessed fMRI, preprocessed T1w, template | aligned fMRI | Figure S7 |
| Nuisance Correction | aligned fMRI | cleaned fMRI | Figure S8 |
| Connectome Estimation | cleaned fMRI, parcellations (Appendix G) | downsampled timeseries, connectome | Figure S9 |

## Appendix C Functional Pipeline

Here we take a deep-dive into each of the modules of the NDMG-f pipeline. We will explain algorithm and parameter choices that were implemented at each step, and the justification for why they were used over alternatives. All QA/QAX figures shown are from participant 0025864 from the BNU1 [21] study.

**Figure S5:**
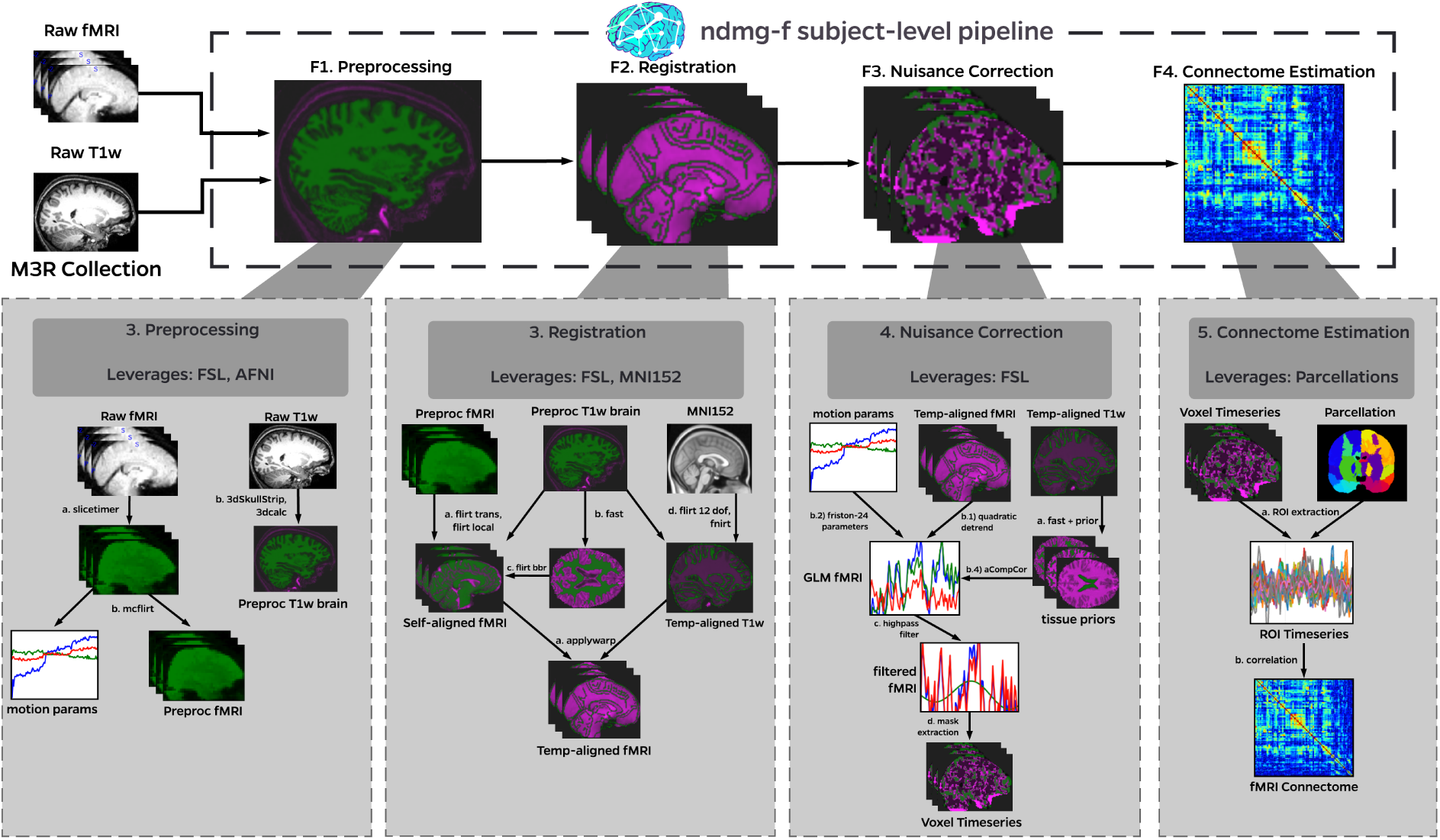
NDMG-f detailed pipeline. The NDMG-f pipeline consists of 4 main steps: **(F1)** Preprocessing, **(F2)** Registration, **(F3)** Nuisance Correction, and **(F4)** Graph Generation. Each of these sections leverages publicly available tools and to robustly produce the desired derivative of each step. Alongside derivative production, NDMG-f produces QA figures at each stage, as can be seen in **F1-F4**, that enable qualitative evaluation of the pipeline’s performance.

### Appendix C.1 Preprocessing

fMRI pre-processing consists of several steps described below (Figure S6).

#### Slice Timing Correction

Slice-timing is accomplished using the slicetimer utility provided by FSL [59]. To collect an individual 4D EPI sequence, a 3D volume is constructed as a combination of individual 2D slices. The 2D slices are collected incrementally; that is, we collect each 2D slice for approximately 10 milliseconds, and the entire 3D volume is complete in about 30 2D slices. This gives a repetition time, or TR, for the volume of seconds, depending on the scanner. The data are essentially a “sliding snapshot” over the course of one TR; observations are not all at a fixed point in time.

NDMG-f accounts for slice timing aliasing by accepting a user-provided acquisition sequence (the order in which slices are collected for a single time point). Given the acquisition sequence of the 2D slices, we can compute the TR shift of each 2D slice using the TR information from the header of the brain image. A slice that occurs first in a TR will have a shift of 0, while a slice that occurs at the end of a TR will have a shift of 1. A slice that occurs exactly in the middle of a TR has a shift of 0.5. For each voxel in an individual slice, interpolation is used to re-center our observations to all have a TR shift of 0.5. For slice-timing correction, NDMG-faccepts a slice-timing configuration file, or one of the canonical slice-timing orientations (interleaved, sequential ascending, or sequential descending).

#### Motion Correction

Motion correction is implemented using the mcflirt utility [58], which is a simplification of FSL’s FLIRT registration tool. During an fMRI scan, participants sit in a small, cramped scanner, often for 5 to 10 minutes. During the course of a study, it is fairly common for participants to move their heads, even if only small amounts. Small shifts will lead to a person’s head being in different spatial positions at each timestep [66], which hampers our efforts to standardize the spatial properties of each individual’s brain down the line through registration. This is because registrations are performed by estimating the registration on the first volume [38–40], after which the estimated transformation is applied across the temporal dimension. This means that if each 3D volume is not aligned spatially, inconsistencies in registration will generally decrease functional connectome quality [86].

Fortunately, given that the individual’s brain structure for each 3D volume is relatively constant (the brain shape itself is not changing in time), a 6 degree of freedom rigid affine transformation for each 3D volume (1 translational and 1 rotational parameter per *x,y,z* direction the individual could move his/her head) using the mean fMRI slice as the reference, adequately addresses this issue.

#### Anatomical Preprocessing

To preprocess the anatomical t1w image, AFN I’s 3dSkullstrip [57] is used. 3dSkull-strip provides modifications to the BET algorithm [87] to make it more robust without hyperparameters. Note that 3dSkullstrip renormalizes intensities, so to regain the original intensities, the result is binarized and fed as a step function (essentially making it a mask) through 3dcalc and multiplied voxelwise with the original image, yielding the original image intensities of the brain and excluding the regions determined to be skull.

### Appendix C.2 Registration

#### Self Registration

To register our input fMRI to our reference T1w image, a three degree of freedom (DOF) affine transformation is estimated with x, y, and z translational parameters with FSL’s FLIRT [88] using the `sch3Dtrans3dof` schedule file provided as part of the FSL package. The schedule file centers the functional brain on the T1w brain and tends to improve the initialization of registrations in later steps. Next, a locally-optimized transformation from the fMRI brain to the T1w brain is estimated. Again, this transformation is robust, and its hyperparameters are tuned to focus on local features of the input (fMRI) and reference (T1w) spaces in the `simple3d.sch` FLIRT schedule file. This schedule file is chosen due to the input fMRI potentially having a narrow field of view, resolution constraints, or tearing that will perform poorly using a more global alignment.

**Figure S6:**
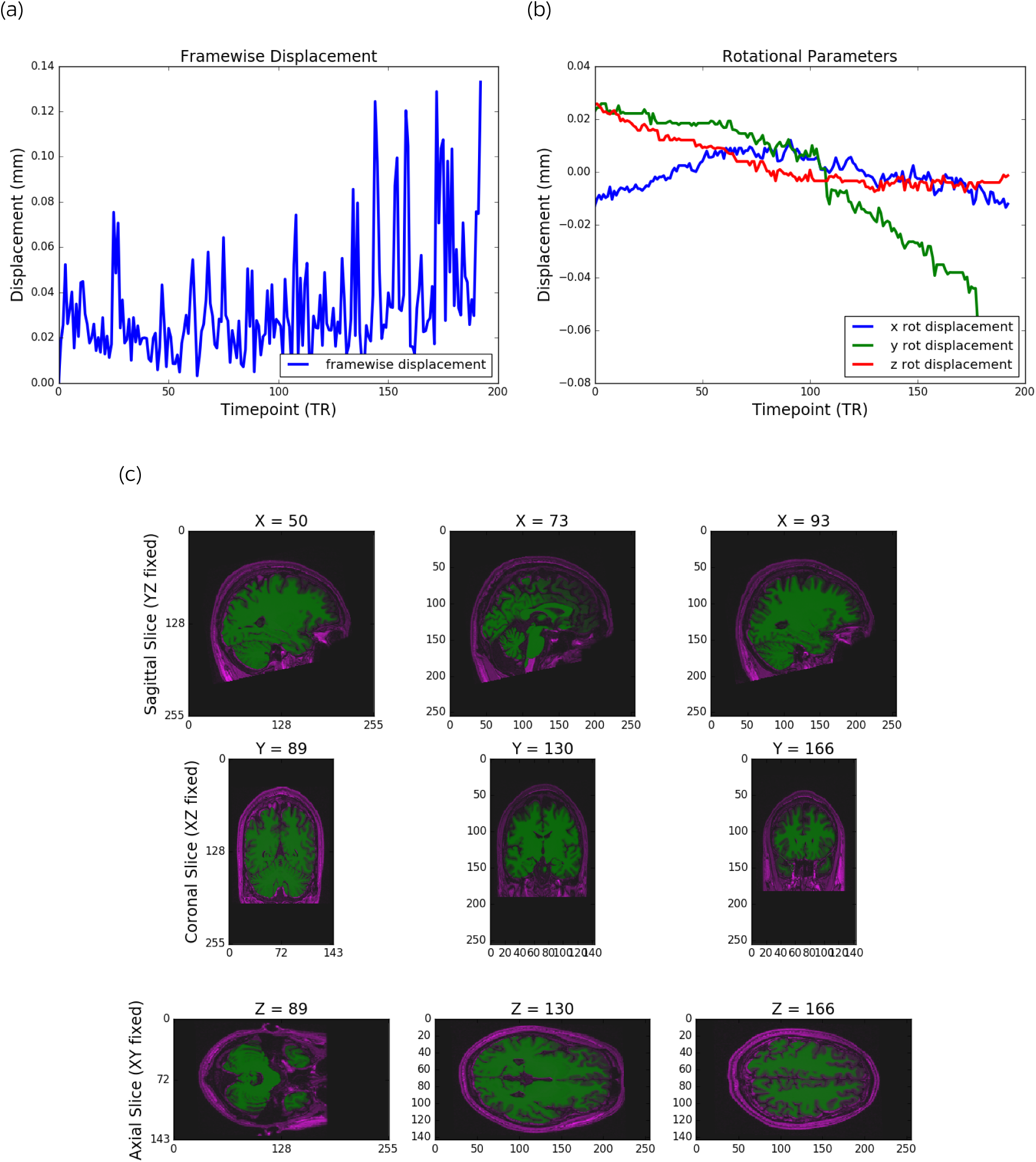
NDMG-f Preprocessing QA. For preprocessing, NDMG-f produces QA figures showing (a) the framewise displacement per timestep, (b) the rotational and translational motion parameters, and (c) plots of the raw and corrected brain, and the success of the brain extraction process.

**Figure S7:**
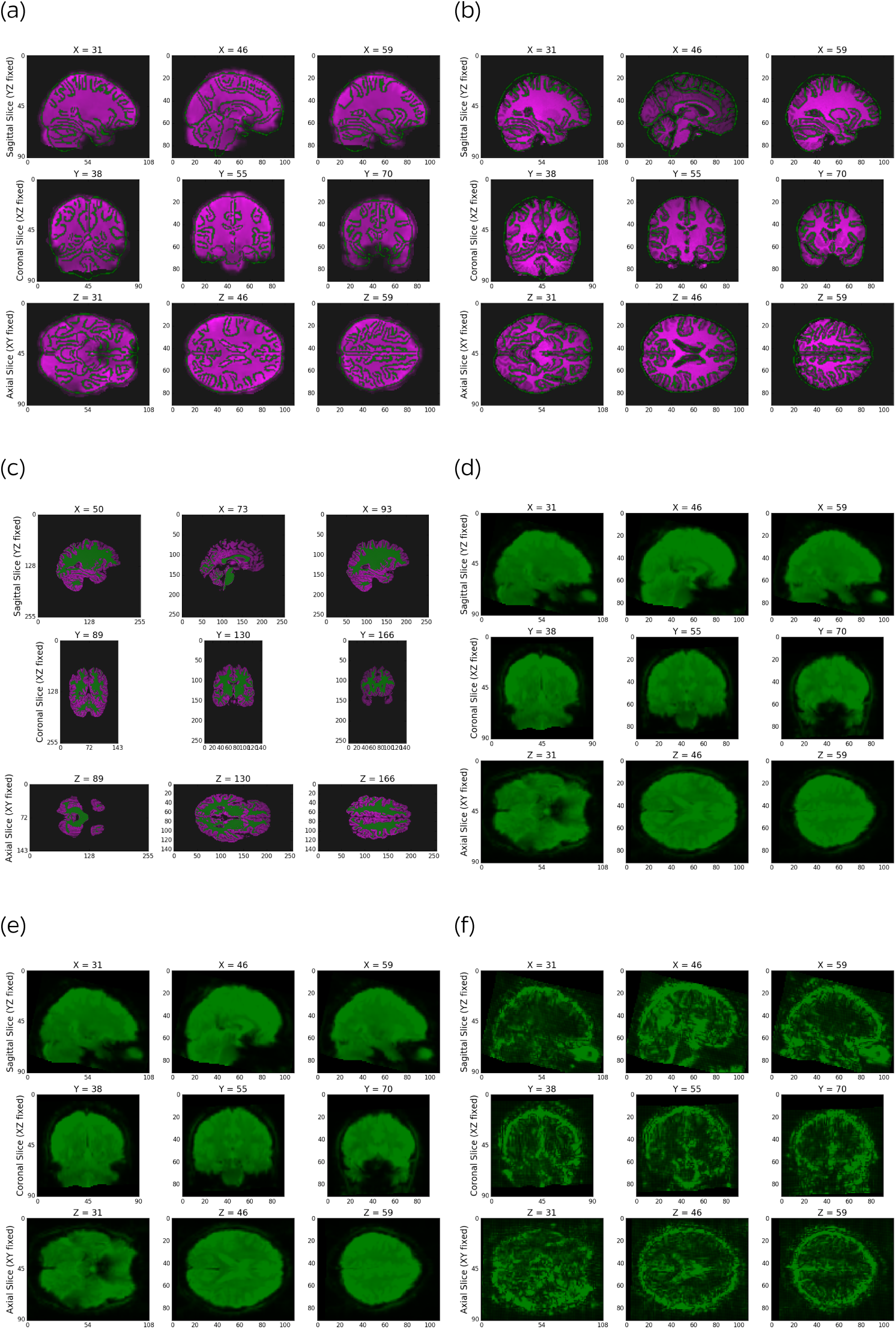
NDMG-f Registration QAX. For registration, NDMG-f produces summary figures showing the preprocessed epi image overlaid on the t1w, the registered epi image overlaid on the template (S7a), the registered t1w image overlaid on the template (S7b), the white-matter mask used in FLIRT-bbr (S7c), the voxelwise mean intensity (3), the voxelwise signal-to-noise ratio (S7e), and the voxelwise contrast-to-noise ratio (S7f).

A third alignment is then estimated using the locally-aligned fMRI to the structural T1w image using the boundary-based registration (`bbr`) cost function provided by FSL [60]. In fMRI, the white-matter/gray-matter border is fairly apparent because the gray-matter generally shows higher intensities than the white-matter regions. Leveraging this observation, the white-matter boundary can be aligned between the fMRI and the T1w scan with high accuracy [60]. A six DOF transformation from the fMRI to the T1w image is estimated, and the T1w is then segmented to produce a white-matter mask using FSL’s `FAST` algorithm [89]. FLIRT is used with the `bbr` cost-function to align the boundaries of the white-matter in the fMRI and T1w scans optimally. This provides a high-quality alignment for intra-modal registration from an EPI to a T1w image.

#### Template Registration

A gentle linear transformation of the T1w brain to the MIN152 template [90] is estimated using the local-optimization schedule file from before. We use this local-optimization registration as the starting point for a more extensive 12-DOF global FLIRT alignment than the self-registration case. Given that the template brain will theoretically be less similar than simple translations, rotations, and scalings can provide, a non-linear registration is estimated from the T1w to the template space. This is accomplished using FSL’s `FNIRT` algorithm [61], with hyper-parameter tuning specific for the MNI152 template. This non-linear transformation is applied first to the T1w image. The non-linear transformation is then combined with the result of the self-alignment step and applied to the functional volume. Applying the transformation only once prevents unnecessary fixed-precision multiplications, which can induce numerical errors.

QA figures for the `registration` step include the preprocessed epi image overlaid on the t1w, the registered epi image overlaid on the template, the registered t1w image overlaid on the template, the white-matter mask used in FLIRT-bbr, the voxelwise mean intensity, the voxelwise signal-to-noise ratio, and the voxelwise contrast-to-noise ratio.

### Appendix C.3 Nuisance Correction

#### General Linear Model

Over the course of an fMRI scanning scan, many sources of noise arise that must be corrected for in order to make quality data inferences. The scanner heats up during a scan (producing a high strength magnetic field for scans lasting up to ten or more minutes, which in turn produces an enormous amount of heat). As the scanner heats, the signal recorded tends to drift (first demonstrated by [63] who showed that a heated scanner detected “brain activity” in cadavers). This drift has been shown to be approximately quadratic [62], so NDMG uses a second-degree polynomial regressor.

While spatial motion correction removes the visual impact of head motion, spurious signal artifacts remain present. These artifacts can be characterized by the position of the brain in the scanner over time [66]. This temporal relationship can be effectively captured by the current volume and the preceding volume, as well as their squares, so 24 regressors are estimated where we have four regressors (current frame, shifted frame, squared-current frame, square-shifted frame) for each of our six (x, y, z, translation and rotation) motion regressors. These regressors are known as the Friston 24 parameter regressors.

Finally, fMRI signal is often corrupted by physiological noise, such as blood flow or vessel dilation [64]. The top 5 principal components from the white-matter and lateral-ventricle signal capture these additional sources of variance [64; 65]. We estimate CSF and white-matter masks using the FAST algorithm, [89] with priors obtained from the MNI152 parcellation [42]. This estimated mask is eroded by 2 voxels on all sides to avoid any potential signal distortion from the gray-matter signal, since gray-matter signal is expected to correlate with our stimuli. Any signal bleeding into the white-matter voxels (since the gray-matter/white-matter boundary has a slight bleed-over region) that could get removed by our PCA could be detrimental to our downstream inferences.

The regressors are incorporated into the design matrix *X* of the general linear model (GLM) shown in (2). For our *n* voxels, the *t* timestep BOLD signal, we can decompose *Y_raw_* ∈ ℝ*^t×n^* as:

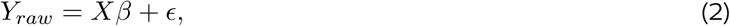

where *ϵ* ∈ ℝ^t×n^ is a noise term, *X* ∈ ℝ^t×r^ is the design matrix, and *β* ∈ ℝ*^t×n^* are the regression coefficients. Minimizing the squared-error loss of *Y_raw_* with respect to *Xβ* will provide an estimate of the coefficients 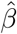 of the regressors in *X:*

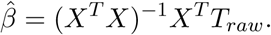

Using the estimate 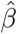 for our regressors *X*, the GLM-corrected timeseries is [62]:

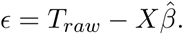

#### Low Frequency Drift Removal

Using the GLM-corrected timeseries, low-frequency drift that may still be present in the functional volume can then be removed. Any physiological response due to a stimulus will have a period of around 16 seconds (or a frequency of 0.063 Hz) and will not exceed the period of any stimuli present, as has been shown in [67]. Using this information, sinusoidal fourier modes with frequencies lower than most brain stimuli are removed. Conservatively, we set a threshold of 0.01 Hz for highpass-filtering out low-frequency noise (this should not remove task-dependent signal as long as our task has a period less than about 100 seconds). Although in this manuscript we only applied NDMG to resting state data, we have also applied it to task data, motivating these thresholds.

#### Tl Effect Removal

During the fMRI scan, the first few volumes may appear to have brighter intensities as the Tl effects are not fully saturated [68]. NDMG therefore discards the first 15 seconds of the fMRI sequence.

### Appendix C.4 Graph Estimation

Given a parcellation with vertices *V* and a corresponding mapping *P*(*v_i_*) indicating the voxels within a region *i*, we first must downsample our voxelwise timeseries. Where *I*(*v*)*_t_* is the intensity at timestep *t* for some *v*. Here, we let 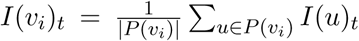, that is, the downsampled timeseries for a given region is the average timeseries over all voxels in that region.

The ROI timeseries can be used to estimate a functional connectome. For a graph 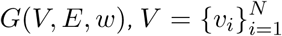 are all of the vertices in the parcellation, *E* = *V* × *V* is all possible edges between vertices, and *w*: *V* × *V* → ℝ, 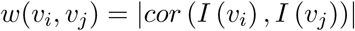 is the absolute temporal correlation of the ROI timeseries between regions *v_i_* and *v_j_*. NDMG computes the pairwise correlation between each pair of regions. The delination of regions is provided by the 24 parcellations described above.

For each parcellation, the correlations are converted to relative ranks—the lowest correlation gets a rank 1, the second lowest rank 2, etc.—and define the ranks as the edge-weights of the resulting functional connectome [33]. QA depicts Figure S9.

## Appendix D Multi-scale Multi-Connectome Analysis

NDMG computes eight node- or edge-wise statistics of each connectome. Each illustrates a non-parametric graph property. The graph statistics are primarily computed with NetworkX and Numpy, and all implementations for NDMG live within the `graph_qa` module. Table 5 provides further information for each statistic.

### Appendix D.1 Group-Level Multi-Scale Analysis

Figure S10 top panel shows the group-level summary statistics of diffusion connectomes belonging to same dataset over 13 parcellations ranging from 48 nodes up to 500 nodes; for clarity, an additional 11 parcellations with up to over 70,000 nodes are not shown here. The bottom panel shows the group-level summary statistics of functional connectomes belonging to the same dataset over 5 parcellations ranging from 52 to 200 nodes. For each parcellation, vertex statistics are normalized by dividing them into number of vertices in the parcellation, and then smoothed via kernel-density estimation to enable comparison across scales. The kernel-width was computed using Scott’s Rule, the default mode for Scipy (https://docs.scipy.org/doc/scipy-0.19.1/reference/generated/scipy.stats,gaussian_kde.html). For most of the statistics, the “shape” of the distributions are relatively similar across scales, though their actual magnitudes can vary somewhat dramatically.

**Figure S9:**
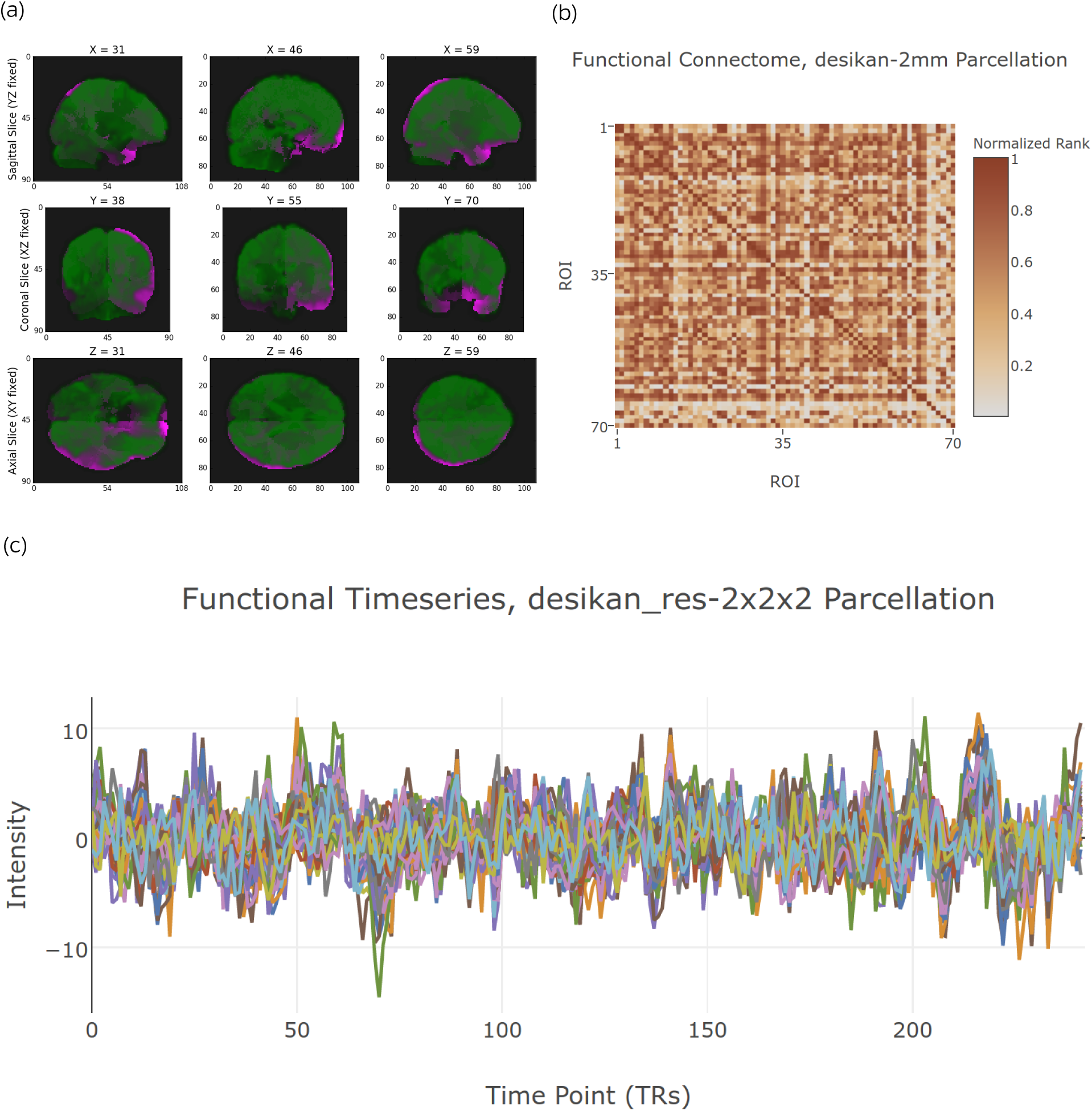
NDMG-f Graph Generation QA. For each parcellation, we visualize (a) the T1w image (green) over the parcellation (purple), the cleaned fMRI over the parcellation, (b) the correlation matrix produced for each parcellated timeseries, and (c) the parcellated timeseries.

**Table 5:** Graph statistics. Each of the graph statistics computed by NDMG. The binarized graphs for NDMG-d were formed by thresholding the non-zero edges. The binarized graphs for NDMG-f were formed by thresholding edges with correlations greater than 0.1, which was identified in [33] as having the highest discriminability for functional connectomes.

| Statistic | NDMG-d | NDMG-f | Implementation |
| --- | --- | --- | --- |
| Betweenness Centrality | Binarized Graph | Binarized Graph | NetworkX |
| Clustering Coefficient | Binarized Graph | Binarized Graph | NetworkX |
| Degree Sequence | Binarized Graph | Weighted Graph | NetworkX |
| Edge Weight Sequence | Binarized Graph | – | NetworkX |
| Eigen Values | Graph Laplacian | Graph Laplacian | NetworkX and Numpy |
| Locality Statistic-1 | Binarized Graph | Weighted Graph | ndmg and NetworkX |
| Number of Non-Zero Edges | Binarized Graph | Binarized Graph | NetworkX |
| Path Length | – | Weighted Graph | NetworkX |
| Cohort Mean Connectome | Weighted Graph | Weighted Graph | Numpy |

### Appendix D.2 Multi-Study Analysis

Figure S11 top panel shows a variety of uni- and multi-variate statistics of the average diffusion connectome from each of the studies enumerated in Table 2 with diffusion data using the Desikan parcellation. The bottom panel shows the same statistics computed on the average functional connectome from each of the studies enumerated in Table 2 with functional data using the Desikan parcellation. In both the diffusion and functional connectomes, each dataset largely appears to have similar trends across each of the statistics shown.

## Appendix E Statistical Connectomics using a Structured Independent Edge Model

Let *g* = (*V, E*) be a graph, where *V* is the set of vertices that is shared for all *i*, and *E_i_* is the set of binary undirected edges between pairs of vertices. Let *A* be a binary adjacency matrix where *a_uv_* = 1 if and only if there is an edge between *u* and *v*, that is, (*uv*) ∈ *E*. Assume *g* is a realization of a random graph *G* ~ *F*, which is sampled from a distribution *F*. We consider a random graph models that generalizes the stochastic block model, the structured independent edge models (SIEM).

The SIEM implies that *A* ~ *SIEM*(*P, τ*), where *τ* is a grouping of the *|E|* edges in *G* into *C* non-overlapping communities, that is, 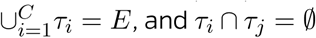 for all *i* ≠ *j*. *P* = [*p*_1_; *p*_2_; *p*_1_; *p*_2_] represents the parameters for within and between edge group probabilities.

Our hypothesis test can be stated as follows:

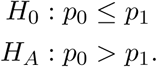

Ignoring the subscript for notational convenience, when all edges in a group are sampled independently with probability *p*, then the number of edges in that group follows a binomial distribution. The maximum likelihood estimate of *p* is 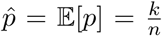, where *k* is the number of observed edges in the group, and *n* is the number of potential edges in the group.

We utilize large sample size theory to obtain a p-value. When *n* is sufficient large, 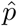 has a normal distribution, 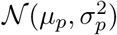 where *μ_p_* = *p* and 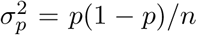. Plugging in 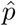 for *p* yields our estimate of the distribution of 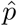. Because 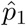 and 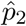 are different, testing whether they are statistically significantly different boils down to testing whether two Gaussians with different variances are different. Welch’s T-Test [91] for testing whether populations have equal means given that they have different variances in the univariate case provides out test statistic:

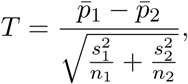

where 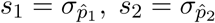 The degrees of freedom can be calculated as follows:

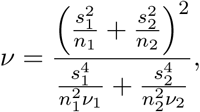

where *v*_1_ = *n*_1_ − 1, *v*_2_ = *n*_2_ − 1. The null distribution, and therefore p-value, is available from R, using the `TDist` family of functions from the `stats` package.

**Figure S10:**
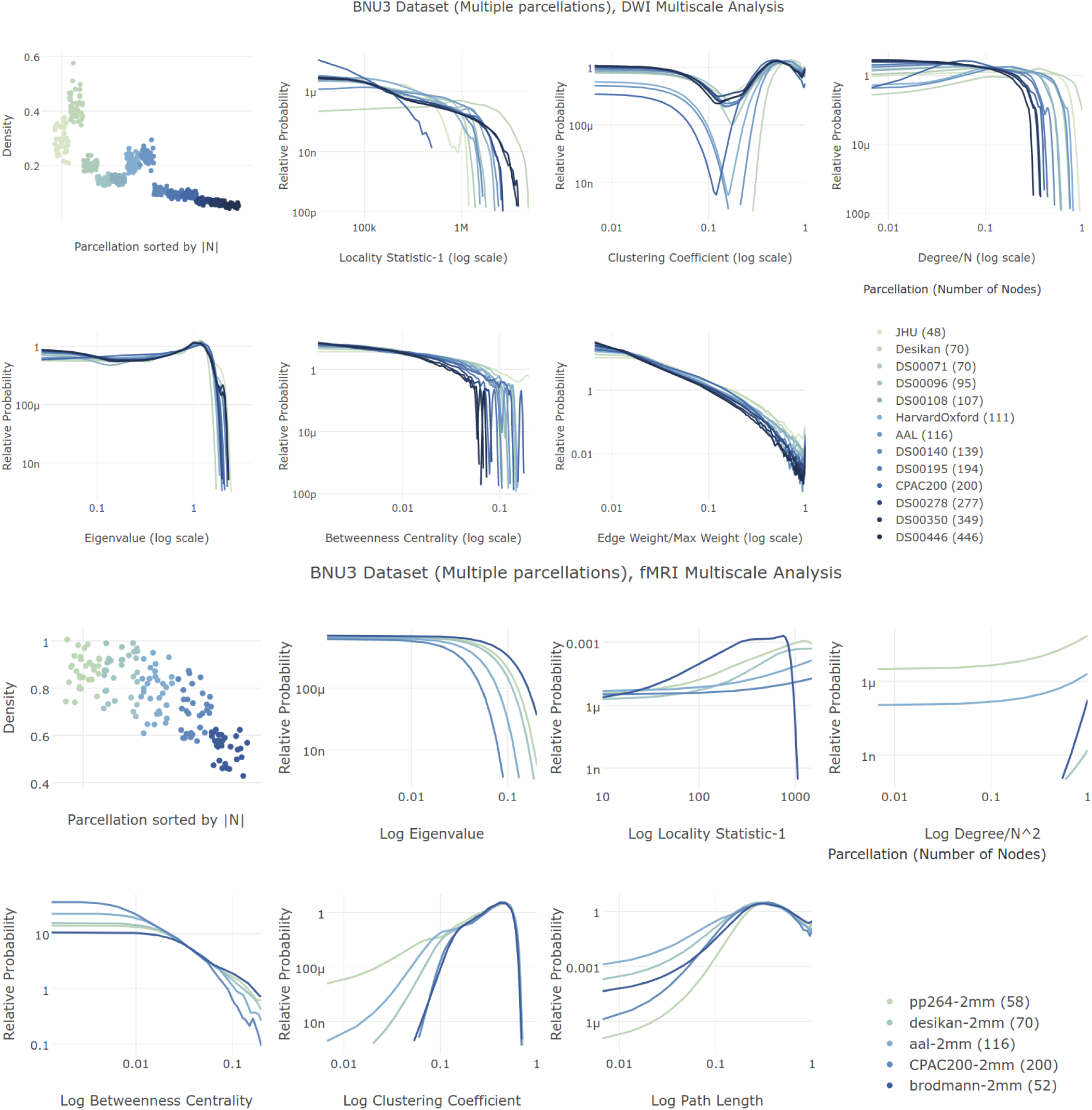
Multi-Scale Connectome Analysis. NDMG produces connectomes at a variety of scales, enabling investigation of graph properties between parcellation schemes. We can observe that the statistics are qualitatively similar in shape across scales, however, they are quantitatively significantly different, for both the diffusion (top) and functional (bottom) connectomes. This suggests that claims made or analyses performed on a given scale may not hold when applied to another scale. This is impactful, as the choice of parcellation has significant bearing on the results of a scientific study.

For the within-modality tests, we first define *p*_1_ as the connection probability within a hemisphere, and *p*_2_ as the connection probability between hemispheres, and *τ* therefore indicates whether a given edge corresponds to an ipsilateral or a contralateral connection. We then redefine *p*_1_ as the connection probability for homotopic edges, and *p*_2_ as the connection probability for heterotopic edges, and *τ* indicates whether an edge corresponds to a homotopic or heterotopic connection.

The across-modality tests use the same basic statistical theory; because everything is Gaussian, linear combinations of Gaussians preserve Gaussianity. Let 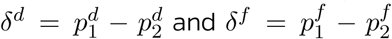. For testing whether the difference between ipsi-lateral and contra-lateral connectivity in the diffusion connectomes exceeds that of the functional connectomes, the across-modality test can be stated as follows:

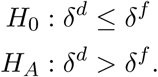

For testing whether the difference between homotopic and heterotopic connectivity in the functional connectomes exceeds that of the diffusion connectomes, the across-modality test can be stated as follows:

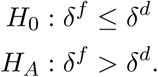

A bit of arithmetic provides the definition of *T* in this setting, again immediately yielding the null distribution and p-value using the `TDist` family of functions from the `stats` package in R.

## Appendix F Batch Effect Investigation

For Batch Effect investigation, we used a simple, intuitive model capturing significant signal as shown in Figure 4. For the dMRI connectomes, our edge communities are the ipsilateral edges in one

community *C_p_* with average ipsilateral (within-hemisphere) community edge weight estimate indicated by 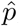, and the contra-lateral (cross-hemisphere) edges *C_q_* with average edge weight estimate 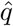. For the fMRI connectomes, our edge communities are the homotopic vs heterotopic. For this investigation, we rank-normalize the edge weights, which has been shown in Wang et al. [33]

to increase the individual-specific signature of brain images. Let *x_i_* = (*p_i_,q_i_*) be the sampled community probabilities for individual *i*, and let *z_i_* denote that individual’s “population” label (described below in more detail). Given a dataset, (*x*_i_, *z_i_*) for *i* ∈ {1,…, *n*}, we assume that each pair is sampled identically and independently from a true but unknown joint distribution: 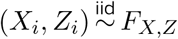 Letting |*z_k_*| denote the number of samples from population *k* yields the following equations for estimating population averaged edge-weights:

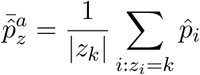

We desire to test whether the population level averages between two populations differ:

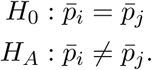

We therefore use the following test statistic: 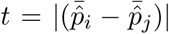, and obtain the null distribution and p-value via a permutation test. To investigate batch effects, we investigate six common pooling strategies with the corresponding labels:

1. Sessions: pooling within a study across unique sessions (Sessions)
2. pooling within a study across unique individuals (Individuals)
3. Sexes: pooling within a study across unique sexes (Sexes)
4. pooling between studies across unique sites (Sites)
5. pooling between unique studies with similar demographics (Demographics)
6. pooling between unique studies with no consideration of demographics (All studies)

## Appendix G Parcellations

**Table 6:**
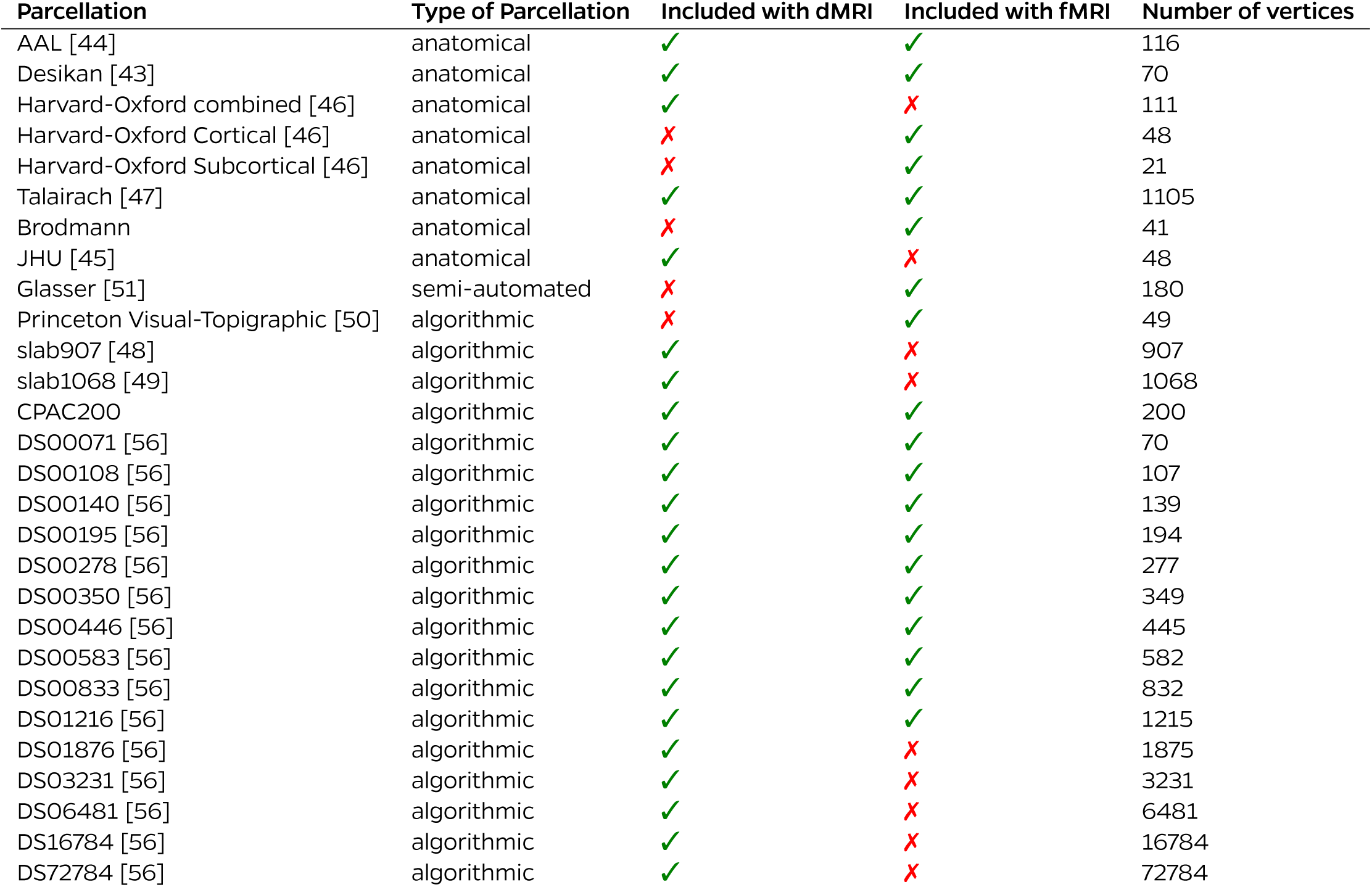
Parcellations. NDMGcomputes graphs at various scales using different parcellation atlases. Parcellations at 1mm were downsampled to 2mm using `flirt` for resampling, with the `nearestneighbour` interpolation method. The number of vertices in the 2mm parcellations used for fMRI may be less than the number of vertices in the 1mm parcellations for the dMRI due to vertices being contracted out by the nearest neighbour interpolation. Below, we distinguish anatomical parcellations, those constructed from anatomical features in the brain, from semi-automated parcellations, those constructed from human-input seeds and adjusted by algorithms, and algorithmic, those constructed purely using algorithms. The parcellations shown below are available for download from our website at `m2g.io`.

| Parcellation | Type of Parcellation | Included with dMRI | Included with fMRI | Number of vertices |
| --- | --- | --- | --- | --- |
| AAL [44] | anatomical | ✓ | ✓ | 116 |
| Desikan [43] | anatomical | ✓ | ✓ | 70 |
| Harvard-Oxford combined [46] | anatomical | ✓ | ✗ | 111 |
| Harvard-Oxford Cortical [46] | anatomical | ✗ | ✓ | 48 |
| Harvard-Oxford Subcortical [46] | anatomical | ✗ | ✓ | 21 |
| Talairach [47] | anatomical | ✓ | ✓ | 1105 |
| Brodmann | anatomical | ✗ | ✓ | 41 |
| JHU [45] | anatomical | ✓ | ✗ | 48 |
| Glasser [51] | semi-automated | ✗ | ✓ | 180 |
| Princeton Visual-Topographic [50] | algorithmic | ✗ | ✓ | 49 |
| slab907 [48] | algorithmic | ✓ | ✗ | 907 |
| slab1068 [49] | algorithmic | ✓ | ✗ | 1068 |
| CPAC200 | algorithmic | ✓ | ✓ | 200 |
| DS00071 [56] | algorithmic | ✓ | ✓ | 70 |
| DS00108 [56] | algorithmic | ✓ | ✓ | 107 |
| DS00140 [56] | algorithmic | ✓ | ✓ | 139 |
| DS00195 [56] | algorithmic | ✓ | ✓ | 194 |
| DS00278 [56] | algorithmic | ✓ | ✓ | 277 |
| DS00350 [56] | algorithmic | ✓ | ✓ | 349 |
| DS00446 [56] | algorithmic | ✓ | ✓ | 445 |
| DS00583 [56] | algorithmic | ✓ | ✓ | 582 |
| DS00833 [56] | algorithmic | ✓ | ✓ | 832 |
| DS01216 [56] | algorithmic | ✓ | ✓ | 1215 |
| DS01876 [56] | algorithmic | ✓ | ✗ | 1875 |
| DS03231 [56] | algorithmic | ✓ | ✗ | 3231 |
| DS06481 [56] | algorithmic | ✓ | ✗ | 6481 |
| DS16784 [56] | algorithmic | ✓ | ✗ | 16784 |
| DS72784 [56] | algorithmic | ✓ | ✗ | 72784 |

**Figure S11:**
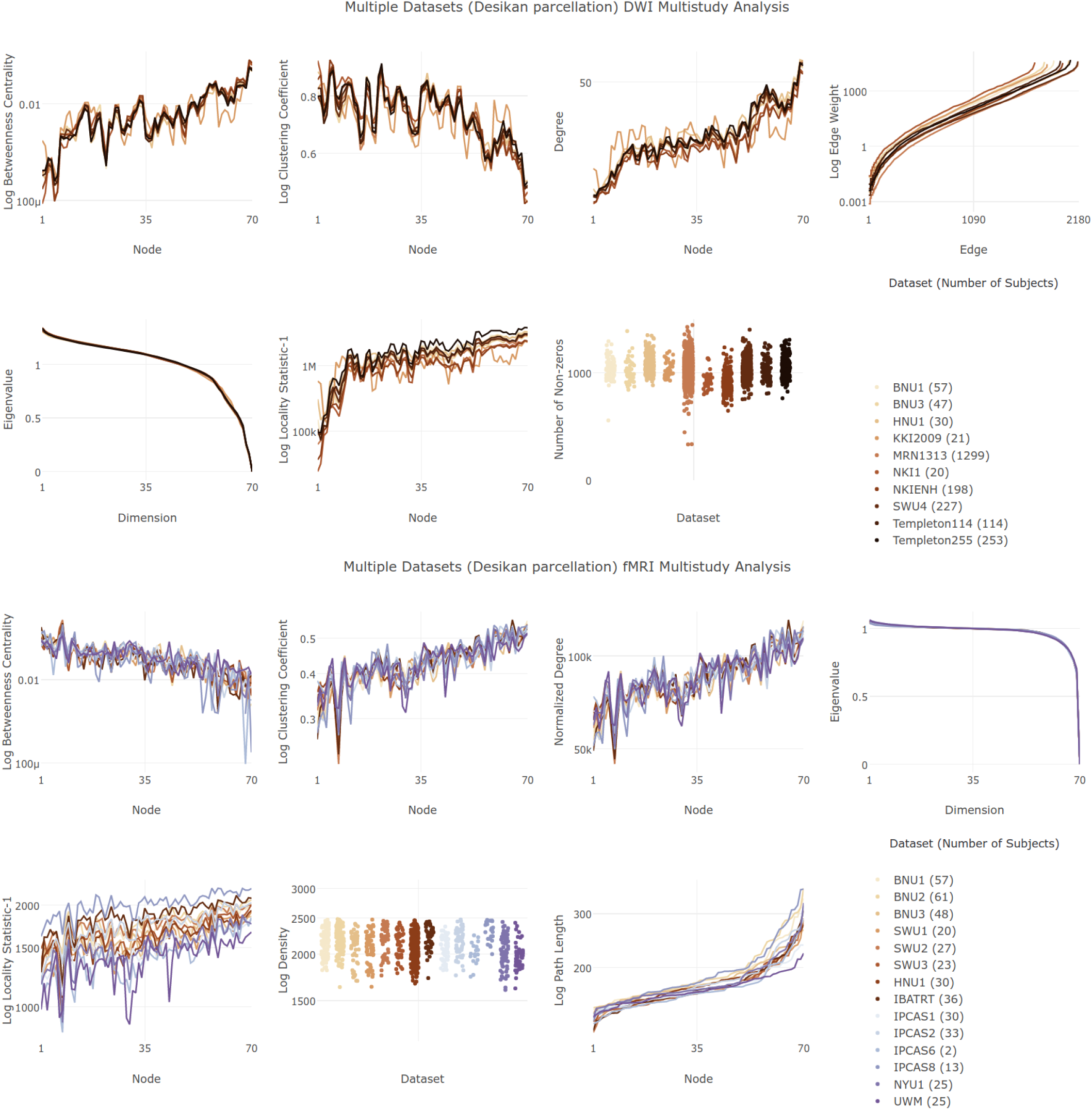
Multi-Study Connectome Analysis. Average connectomes from ten diffusion studies processed with NDMG-d are qualitatively compared by way of their summary statistics on the Desikan parcellation in the top figure. The Desikan atlas used in NDMG has been modified to include two additional regions, one per hemisphere, which fills in a hole in the parcellation near the corpus callosum. The nodes in this plot have been sorted such that the degree sequence of the left hemisphere (Desikan nodes 1–35) of the BNU1 dataset is monotonically non-decreasing, and that corresponding left-right nodes are next to one another. Each line shows the average for each statistic over all individuals within the study. On the bottom, we repeat the same analysis on the functional connectomes from seventeen different studies. Like the statistics computed in for the diffusion connectomes, the statistics are again qualitatively similar but quantitatively disparate. This suggests that claims made or analyses performed on a given scale may not hold when applied to another scale. Again, we see that parcellation choice has an impact on the implications of a study. Information on the graph statistics computed can be found in Appendix D.

